# AGTR1 is overexpressed in neuroendocrine neoplasms, regulates secretion and may serve as a target for molecular imaging and therapy

**DOI:** 10.1101/853788

**Authors:** Samantha Exner, Claudia Schuldt, Sachindra Sachindra, Jing Du, Isabelle Heing-Becker, Kai Licha, Bertram Wiedenmann, Carsten Grötzinger

## Abstract

Peptide receptor targeting has proven to be a pivotal tool for diagnostic imaging and radioligand therapy of neuroendocrine neoplasms (NENs), which frequently express somatostatin receptors (SSTRs) on their cell surface. However, up to 30 % of NEN patients do not benefit from SSTR-based approaches, others develop a resistance. Consequently, alternative cell surface targets need to be identified. In this study, cell-based dynamic mass redistribution and calcium mobilization screening using a 998-compound library identified and confirmed angiotensin II (ATII) as a strong activator of cellular signaling in NEN cells. Expression analyses of the ATII receptor type 1 (AGTR1) revealed an upregulation of both mRNA levels (RT-qPCR) and radioligand binding (autoradiography) in pancreatic (n=42) and small-intestinal (n=71) NEN tissues compared to healthy controls (n=25). The two NEN cell lines BON (pancreas) and H727 (lung) with elevated AGTR1 expression exhibited concentration-dependent calcium mobilization and chromogranin A secretion upon stimulation with ATII, blocked by AGTR1 antagonism and G_αq_ inhibition. To assess the applicability of AGTR1 for optical in vivo imaging, the receptor ligand saralasin was coupled to the near-infrared dye indotricarbocyanine and tested for its biodistribution in a NMRI Foxn1^nu^/Foxn1^nu^ mouse model bearing AGTR1-positive BON and negative QGP-1 xenograft tumors. Near-infrared fluorescent imaging showed a significantly higher uptake in BON tumors 3-6 hours after injection. This successful targeting in an NEN model establishes AGTR1 as an interesting target in this tumor entity, paving the way for the development of translational chelator-based probes for diagnostic PET imaging and peptide receptor radioligand therapy.

## Introduction

Neuroendocrine neoplasms (NENs) represent a heterogeneous group of neoplasms with a moderate but steadily increasing annual age-adjusted incidence of approximately 7/100,000 for the USA in 2012 (1). Most NENs grow rather slowly and may develop into a more aggressive phenotype while being undetected. Tumor growth often goes unnoticed and diagnosis may be delayed for many years after the onset of disease, until metastatic spread. At this point, local surgery as the only curative therapy is not feasible anymore (2). Among treatment options for advanced tumors are targeted therapies with mTOR inhibitors (everolimus), tyrosine kinase inhibitors (sunitinib) and somatostatin analogs (SSAs) as NENs frequently express somatostatin receptors (SSTRs) (3). Autoradiographic receptor detection, in-situ hybridization and immunohistochemistry demonstrated high SSTR2 overexpression in tumors with little background in other tissues (4–6). SSAs such as octreotide and lanreotide are used as antisecretory and antiproliferative medication. Apart from direct pharmacological intervention, SSTR overexpression in NENs has also been utilized for targeted molecular imaging and peptide receptor radiotherapy (PRRT) (7–10).

Although SSTR-based imaging and therapy is an important option in the management of NENs, it is estimated that up to 30 % of patients do not fully benefit from these approaches. Whereas some tumors lack SSTR expression or downregulate expression upon treatment, others exhibit an inhomogeneous receptor distribution resulting in a residual tumor mass after treatment and subsequent resistance to SSAs (11, 12). Other possible resistance mechanisms include: receptor desensitization, downregulation or loss-of-function as well as altered signaling pathways (12, 13). Most patients develop SSA resistance within weeks to months of treatment, with recurrent symptoms and tumor growth (14). As a consequence, to extend the repertoire of available ligands for imaging and therapy, alternative peptide GPCRs are investigated.

To identify alternative target receptors at the cell surface of NENs, this study used high-throughput screening technology to detect G protein-coupled receptor (GPCR) activation in human NEN cells. The screening revealed a clear response towards angiotensin II (ATII). Therefore, this study went on to assess the potential of AGTR1 as a novel target in NEN. Study objectives were to validate AGTR1 mRNA expression and ligand binding in human NEN tissue, to investigate the biological effects of ATII in human cell models and to evaluate the suitability of AGTR1 as a target for tumor diagnosis in a mouse xenograft model.

## Material and Methods Reagents

If not indicated otherwise, chemicals were obtained from Sigma Aldrich (St. Louis, MO, USA). G_αq_ inhibitor UBO-QIC was purchased from the Institute of Pharmaceutical Biology, University of Bonn (Germany), AKT-Inhibitor AZD-5363, #A9601, from LKT laboratories (St Paul, MN, USA), ERK-Inhibitor SCH772984, #B1682-5, from BioVision (Milpitas, CA, USA), β-arrestin inhibitor barbadin, #Axon2774, from Axon Medchem (Groningen, Netherlands).

### Compound library

The compound library used for screening consisted of the following components: compounds 1-866 were from the G-Protein Coupled Receptor (GPCR) Peptide Ligand Library (Phoenix Pharmaceuticals, Burlingame, CA, USA), compounds 867-956 from the Orphan Peptide Ligand Library (Qventas Labs, Branford, CT, USA), and compounds 957-998 from various sources. All compounds in the library are listed in Supplementary Table 1.

### Cell culture

If not indicated otherwise, cell culture reagents were obtained from Biochrom AG (Berlin, Germany). The human NEN cell lines BON and QGP-1 (pancreatic), LCC-18 (colonic), H727 and UMC-11 (pulmonary) and all other cell lines were cultured in RPMI 1640 supplemented with 10 % fetal calf serum in a humidified atmosphere at 37 °C and 5 % CO_2_. BON cells were a kind gift of Courtney Townsend (University of Texas, Galveston, USA). LCC-18 were kindly donated by Kjell Öberg (University of Uppsala, Sweden) and KRJ-1 by Dr. Irvin Modlin (Yale University, USA). QGP-1 were obtained from the Japanese Collection of Research Bioresources. All other cell lines were obtained from Cell Line Services (Heidelberg, Germany), DSMZ (Braunschweig, Germany) or ATCC (Manassas, VA, USA). Cells were cultured for no more than 20 passages. Mycoplasma testing was done at least every six months. Cell lines were authenticated by STR analysis at DSMZ (Braunschweig, Germany). Due to the lack of reference profiles for BON and LCC-18, their respective STR profile was characterized as unique and not contaminated with any known cell line. The results of the STR profiling for BON and LCC-18 are provided in the table below.

**Table 1:** STR profiling data for BON and QGP-1 cells.

|  | D5 | D5' | D13 | D13' | D7 | D7' | D16 | D16' | vWA | vWA' | TH01 | TH01' | TPOX | TPOX' | CSF1 | CSF1' | Amel | Amel' |
| --- | --- | --- | --- | --- | --- | --- | --- | --- | --- | --- | --- | --- | --- | --- | --- | --- | --- | --- |
| **BON** | 9 | 12 | 11 | 12 | 9 | 9 | 10 | 11 | 18 | 19 | 8.0 | 8.0 | 9.0 | 9.0 | 10 | 11 | X | Y |
| **LCC-18** | 14 | 14 | 8 | 11 | 11 | 12 | 10 | 13 | 16 | 19 | 9.0 | 9.3 | 8.0 | 10.0 | 11 | 11 | X | X |

### Tissues

This study was carried out in accordance with the recommendations and protocol approval by the local ethics committee at Charité - Universitätsmedizin Berlin with written informed consent from all subjects, in accordance with the Declaration of Helsinki. Samples were flash-frozen after surgical resection and stored at −80 °C until further use.

### Reverse transcription quantitative real-time PCR (RT-qPCR)

RNA isolation, reverse transcription and quantitative real-time PCR were performed as previously described (15). Oligonucleotide primer sequences are indicated in Table 2. Plotted values were corrected for different primer efficiencies and normalized on the geometric mean of UBC, ALG9 and HPRT1 using the ΔΔCt method (16). Reference genes were validated and chosen using the geNorm algorithm (17) by qbase+ software (Biogazelle, Ghent, Belgium).

**Table 2:**
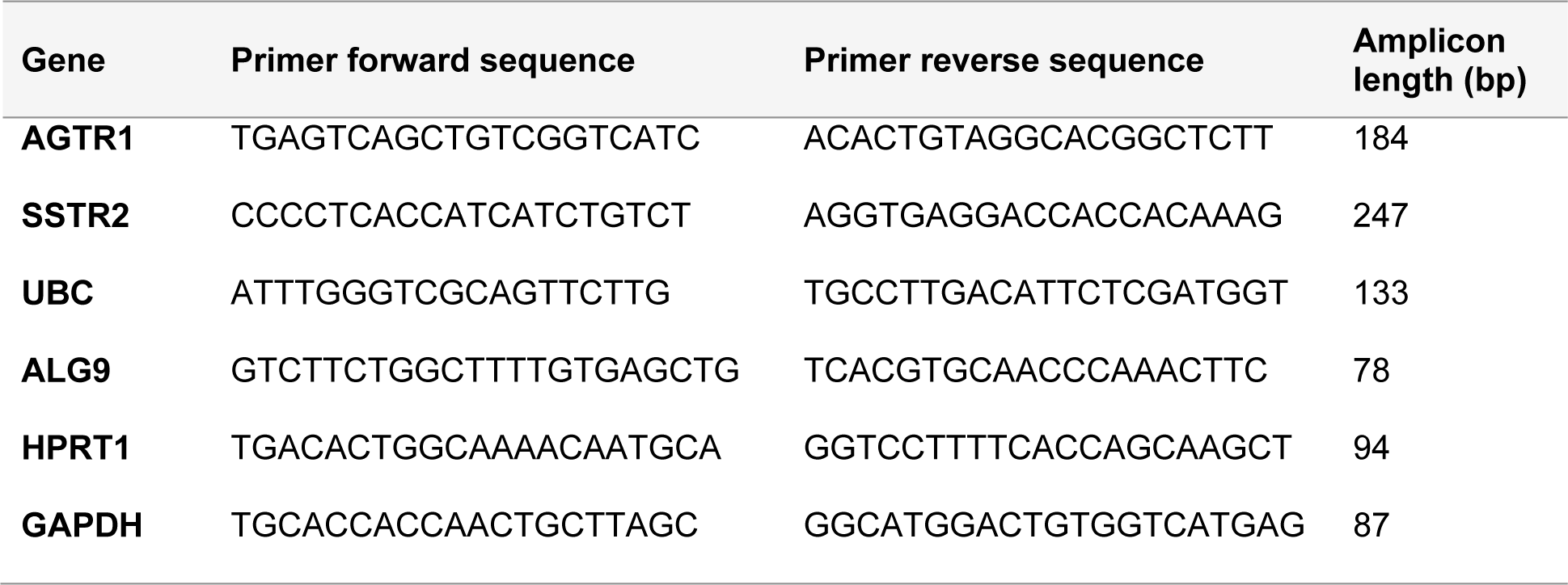
Primer sequences used for RT-qPCR.

### Radioactive studies

Peptide iodination was performed as previously described (18). Briefly, 10 nmol angiotensin II (Sigma, Deisenhofen, Germany) were iodinated with 1 mCi carrier-free Na^125^I (NEZ033L010MC, Perkin Elmer, Waltham, MA, USA) and purified by HPLC (Analytic HPLC 1200 Series, Agilent, Santa Clara, CA, USA; see Figure S1 in supplementary material). The method for the competitive binding assay was described in detail before (15). For saturation experiments, binding buffer was prepared with varying concentrations of ^125^I-angiotensin II, either with (non-specific binding) or without 1 µM of additional unlabeled peptide (total binding). Non-specific binding was subtracted from total binding to obtain specific binding. Obtained cpm values were plotted with GraphPad Prism 5.04 and fitted using nonlinear regression (one site-fit Ki for competition data, one site - total and non-specific/one site - specific binding for saturation data).

### In vitro receptor autoradiography

Flash-frozen human and mouse tissue samples were cut into 10-20 µm thick sections using a cryomicrotome, mounted on glass slides, dried and stored at −80 °C. On the day of the experiment, slides were thawed, tissues encircled with a Dako Pen and incubated with 200 µl binding buffer (50 mM Hepes pH 7.4, 5 mM MgCl2, 1 mM CaCl2, 0.5 % BSA, cOmplete protease inhibitors) containing 0.5 nM radiolabeled ^125^I-angiotensin II (total binding). Non-specific binding was assessed on consecutive sections by additional incubation with 1 µM of unlabeled peptide. After 1-2 h at 37 °C, slides were transferred to glass cuvettes, washed three times for 1 min with washing buffer (50 mM Tris-HCl, pH 7.4, 125 mM NaCl, 0.05 % BSA), shortly dipped into water and quickly dried under a stream of air. Slides were exposed to Amersham Hyperfilm MP (GE Healthcare, Buckinghamshire, UK) for 1-2 weeks at room temperature. Developed films were scanned and analyzed with ImageJ. In addition to densitometric quantification, tissue sections were carefully wiped off the slides using filter paper and measured in a gamma counter. Additional tissue sections were stained with hematoxylin and eosin for comparison with the corresponding autoradiograms.

### Calcium mobilization

Experiments in 96-well format were performed as previously described (18). For 384-well experiments, 15,000 to 20,000 cells per well were seeded after plate coating and for any incubation and washing steps 20 µl total volume were used. For experiments using antagonists, these were diluted in washing buffer/0.5% BSA, added to the cells after the second washing step in double concentration and incubated for 15 min before measurement.

### DMR assay

384-well or 96-well BIND CA1 biosensor plates (SRU Biosystems, Woburn, MA, USA) were seeded with 15,000/50,000 cells per well. The next day, cells were incubated with 25/100 µl serum-free medium for 10 min at room temperature. A buffer baseline was obtained on the dynamic mass redistribution (DMR) reader (BIND reader, SRU Biosystems) for approximately 5 min before compounds were added by a 96-well pipette (Liquidator 96, Mettler Toldedo, Gießen, Germany) within 30-60 sec. DMR data were acquired for 45 minutes and exported using EMS Export Wizard (SRU Biosystems). A compound was regarded a hit if its trace contained at least three data points with a read higher than the mean ± 3 standard deviations of the negative control (buffer only). For EC50 calculations, data points with the highest shift between buffer control and compound dilution were used and plotted with GraphPad Prism 5.04.

### Impedance assay

50,000 BON cells were seeded per well into 96-well impedance sensor plates (Bionas, Rostock, Germany). The next day, cells were incubated with 100 µl serum-free medium for 10 min at room temperature. A buffer baseline was obtained on the impedance reader (Adcon reader, Bionas) for approximately 5 min before compound dilutions (100 µl) were added by a 12-channel pipette within 30-60 sec. Impedance data were acquired for 45 minutes and data points with the highest shift between buffer control and compound dilution were used to calculate EC50 values using GraphPad Prism 5.04.

### Chromogranin A ELISA

30,000 BON cells per well were seeded in 96-well plates and grown overnight. The next day, ligands were diluted in serum-free medium/0.5 % BSA and the used medium was aspirated before adding 250 µl of the respective dilution to each well. Cells were incubated for 6 h at 37 °C. Supernatants were then transferred into a U-bottom plate, centrifuged for 3 min at 800 x *g* and carefully transferred into another U-bottom plate. Supernatants were directly used for ELISA measurements according to the manufacturer’s protocol (Chromogranin A ELISA Kit, Dako, Glostrup, DK). Inhibitor experiments were performed in complete medium for 24 h after ligand application and supernatants were processed as described.

### Metabolic activity

Cells were seeded in quadruplicates at a density of 5,000 cells in 50 µl medium per well in 96-well plates, grown overnight and treated with the indicated ligand concentrations (on top of the well, final volume 100 µl). After 96 h, metabolic activity was determined by addition of 100 µl medium containing AlamarBlue™ redox indicator (Thermo Fisher, Waltham, MA, US) to each well. After 3-4 h, the resulting fluorescence was measured with an EnVision Multilabel Plate Reader (Perkin Elmer, Waltham, USA). Values were normalized as percent of control treated with vehicle.

### AGTR1 ligands and ligand conjugates

Angiotensin II as well as saralasin-ITCC (the dye indotricarbocyanine [ITCC] linked via 4,7,10-trioxatridecan-succinamic acid [TTDS]) were obtained from peptides&elephants (Hennigsdorf, Germany). Valsartan and PD102807 were from Tocris (Bristol, UK). Valsartan-ITCC (the dye ITCC linked via 1,3-diamino propane) was synthesized as described in the supplementary methods.

### In vivo imaging

8-week-old immunodeficient NMRI-*Foxn1^nu^/Foxn1^nu^* mice (Janvier Labs, Le Genest-Saint-Isle, France) were inoculated with BON and QGP-1 cells to generate xenografts for in vivo experiments. Tumor cells grown as monolayers in culture plates were harvested, counted and the pellet was resuspended in serum-free medium at 1 x 10^7^ cells/100 µl. Cells were mixed 1:1 with Matrigel HC (Corning, Corning, USA), subcutaneously injected into the shoulder of anesthetized mice (100 µl, 5 x 10^6^ cells) and grown for 2-3 weeks. For in vivo receptor targeting, the near-infrared fluorescent ITCC-labeled probe was applied intravenously into tumor bearing mice via a lateral tail vein (1 nmol in 100 µl 0.9 % NaCl per animal). Images were acquired prior to and at the indicated time points after injection with a Pearl near-infrared imaging system (Licor, Lincoln, USA). Animals were imaged under isoflurane anesthesia with an excitation at 785 nm and an emission at 820 nm. Images were analyzed with the Pearl Cam Software by defining regions of interest to obtain mean fluorescence intensities. From these, the ratio of signal (tumor) to background (neck) was calculated. For ex vivo imaging, mice were sacrificed at 6 h after injection and tumors and organs were surgically excised. Animal care followed institutional guidelines and all experiments were approved by local animal research authorities.

## Data availability

Numerical data for all applicable experiments (xlsx file) have been deposited in an open data repository for public access: https://doi.org/10.5281/zenodo.3550904

## Results

In order to identify potential cell surface targets in NEN cells, a 998-compound chemical library mainly consisting of peptide receptor ligands (Supplementary Table 1) was applied in a primary screening for cellular activity in the human pancreatic NEN cell line BON. Both a label-free dynamic mass redistribution (DMR) assay as well as an intracellular calcium mobilization assay were employed in 384-well format (Fig. 1A-C). A total of 89 hits were detected (8.9 %), which were assayed again in BON cells in a 96-well format (Fig. 1D-E). 30 primary hits, including structurally related compounds, were confirmed in 96-well format in the DMR assay, and 19 hits in the calcium assay, with 11 compounds confirmed in both assays (Supplementary Table 2). Angiotensin II (ATII) appeared consistently as a strong activator of intracellular signaling in BON cells. This was subsequently validated in concentration-response experiments using these cells in three different functional assay types: DMR, calcium and impedance. The quantitative analysis showed an EC50 for ATII of 0.88 ± 0.28 nM, 4.9 ± 5.2 nM and 0.45 ± 0.03 nM in DMR, calcium and impedance measurements, respectively (Fig. 1F-H).

**Figure 1:**
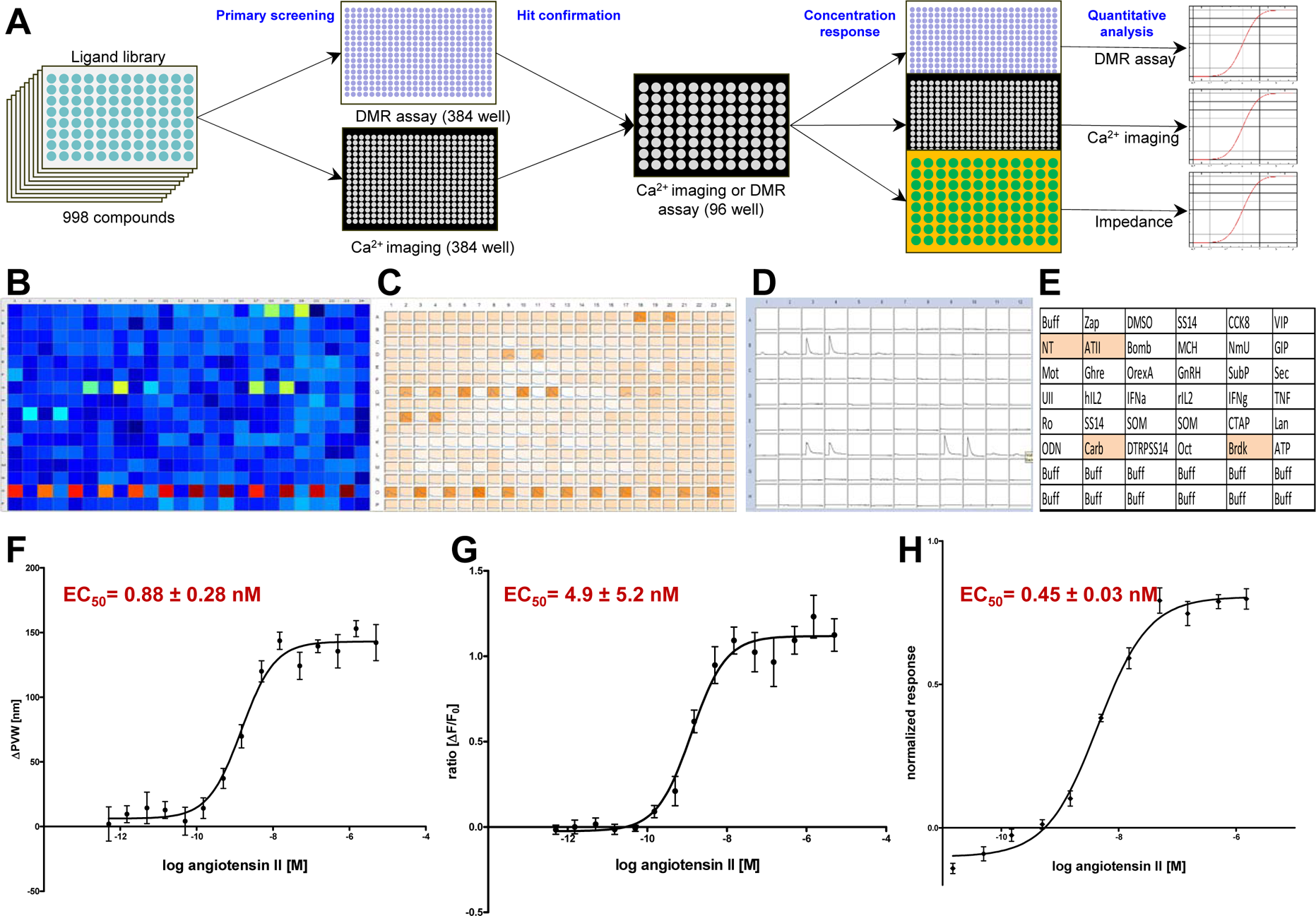
Identification and confirmation of ATII-induced signaling in NEN cells. **(A)** Schematic workflow of the screening and hit confirmation approach. A 998-compound library of peptides and small molecules was used in the primary screening on neuroendocrine BON cells, applying both DMR and calcium mobilization assay in the 384-well format, followed by hit confirmation using the calcium assay (96-well format). Finally, DMR, calcium and impedance assays were used to determine quantitative pharmacological data. **(B)** Pseudocolor representation of the screening result from one DMR assay plate. Duplicates are interspaced by one well, penultimate row contains positive and negative controls. **(C)** Representative results from calcium assay, plate layout as in (B). **(D)** Plate view of hit confirmation calcium assay with fluorescence intensity traces for each well. **(E)** Plate layout with ligands used for hit confirmation, plate as in (D). **(F-H)** Concentration-response curve for angiotensin II in BON cells measured by DMR assay (F), calcium mobilization (G) and impedance (H).

After functional detection of ATII-induced signaling in a human NEN cell line, endogenous expression of the ATII receptor AGTR1 was assessed in five NEN cell lines by RT-qPCR and compared to mRNA levels of 23 other tumor and non-tumor cell lines (Fig. 2A). BON and H727, originally isolated from NENs of pancreas and lung, exhibited the highest mRNA levels of all tested cell lines, indicating a potential AGTR1 overexpression in NEN. NEN cell lines QGP-1 (pancreas) and LCC-18 (colon) showed at least 1000-fold lower mRNA expression and were employed as AGTR1-negative NEN control cell lines. UMC-11 cells (lung) showed intermediate AGTR1 mRNA abundance.

**Figure 2:**
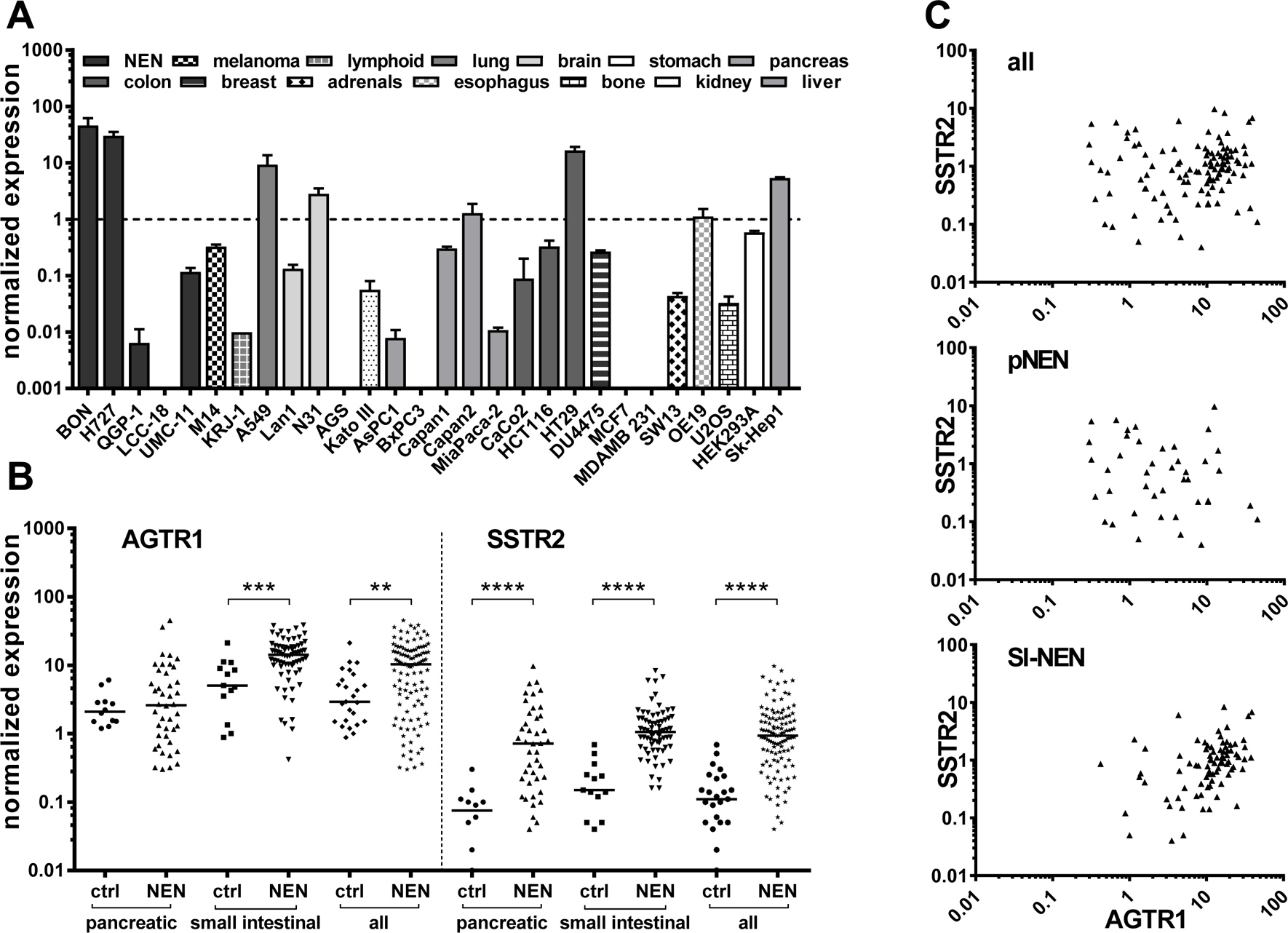
Increased AGTR1 gene expression in pancreatic and small intestinal NEN tissue and two NEN cell lines. **(A)** AGTR1 expression was measured as mRNA levels by RT-qPCR in 28 cell lines of different origin. Values were normalized on UBC, HPRT1 and GAPDH. Bars show mean ± S.E.M (n=2, different passages). **(B)** AGTR1 mRNA levels determined by RT-qPCR in pancreatic (n=42) and small intestinal (n=71) NEN tissues in comparison to pancreatic (n=12) and small intestinal (n=13) control tissues. Values were normalized on ALG9, UBC and HPRT1 and compared to SSTR2 gene expression in the same samples. Bars represent median. Evaluated with unpaired two-tailed Student’s t-test, **p<0.01, ***p<0.001, ****p<0.0001. **(C)** Scatter plots showing the correlation of AGTR1 with SSTR2 gene expression in pancreatic (pNEN) and small intestinal (SI-NEN) tumor tissues.

To further investigate AGTR1 expression in this tumor entity, pancreatic (n=42) and small-intestinal (n=71) tumor tissues of NEN patients as well as control tissues (n=25) were analyzed for their AGTR1 mRNA levels. Small-intestinal NEN (siNEN) demonstrated significantly (p<0.001) increased AGTR1 transcript levels, with an overall 2.8-fold higher median value in comparison to healthy control tissues, whereas pancreatic tissues showed no significant difference (Fig. 2B). Taking the median value of the siNEN samples as a cut-off, 50% of tumor samples yet only 7.7% of controls were above that value. When all NENs were compared to all controls, differences were also significant (3.6-fold higher median, p<0.01). The same samples were also analyzed for expression of the established NEN target SSTR2, which was found to be significantly upregulated in both pancreatic and siNEN tissues, as expected. The ratio of NEN to control median values, was higher for SSTR2 (8.5 versus 3.6 for AGTR1). However, AGTR1 was detected at an approximately 10-fold higher expression level than SSTR2. Correlation analysis confirmed a significant positive correlation between AGTR1 and SSTR2 expression for small-intestinal NENs (Pearson r=0.37, p<0.001, r²=0.14), whereas no significant correlation was found for pancreatic samples (r=-0.09) or all NENs taken together (r=0.12) (Fig. 2C).

To verify AGTR1 gene expression employing an alternative methodology, ten normal control samples from pancreas and small intestine as well as eight pancreatic and nine small-intestinal NEN samples were selected for the determination of receptor binding sites by in vitro receptor autoradiography (patient characteristics: Supplementary Table 3). To this end, the natural receptor ligand ATII was radioactively labeled with iodine-125 (^125^I-ATII) and purified by high-performance liquid chromatography (HPLC) (Supplementary Fig. 1). Consecutive cryosections of each tissue were incubated with the radioligand in the absence or presence of an excess of non-labeled ATII. To distinguish between receptor subtypes, additional tissue sections were incubated with radioligand and AGTR1-specific antagonist valsartan or AGTR2-specific antagonist PD123319 (Fig. 3A, Supplementary Fig. 3-4). Pancreatic NEN tissues showed a rather weak overall binding of ^125^I-ATII. Nevertheless, in tissues #4 (Fig. 3A), #1, #6 and #8 (Supplementary Fig. 3), it could be clearly displaced by ATII and valsartan, but not by PD123319, indicating dominant AGTR1 expression. Small-intestinal NENs, on the other hand, demonstrated a strong binding in more than half of the samples (#9 to #13) (Fig. 3A, Supplementary Fig.3). This proved to be AGTR1-specific binding as it could only be displaced by valsartan. PD123319 was unable to compete with the radioligand. In addition, pancreatic and small-intestinal control tissues were included in the analysis (Fig. 3A, Supplementary Fig. 4). Interestingly, a number of pancreatic samples exhibited strong radioligand binding. In comparison to pancreatic NEN samples, signals in normal pancreas were displaced solely by PD123319, indicating a specific expression of AGTR2. Small-intestinal control samples showed no binding at all (#23, #25) or non-specific binding (#24, #26, #27) (Supplementary Fig. 4).

**Figure 3:**
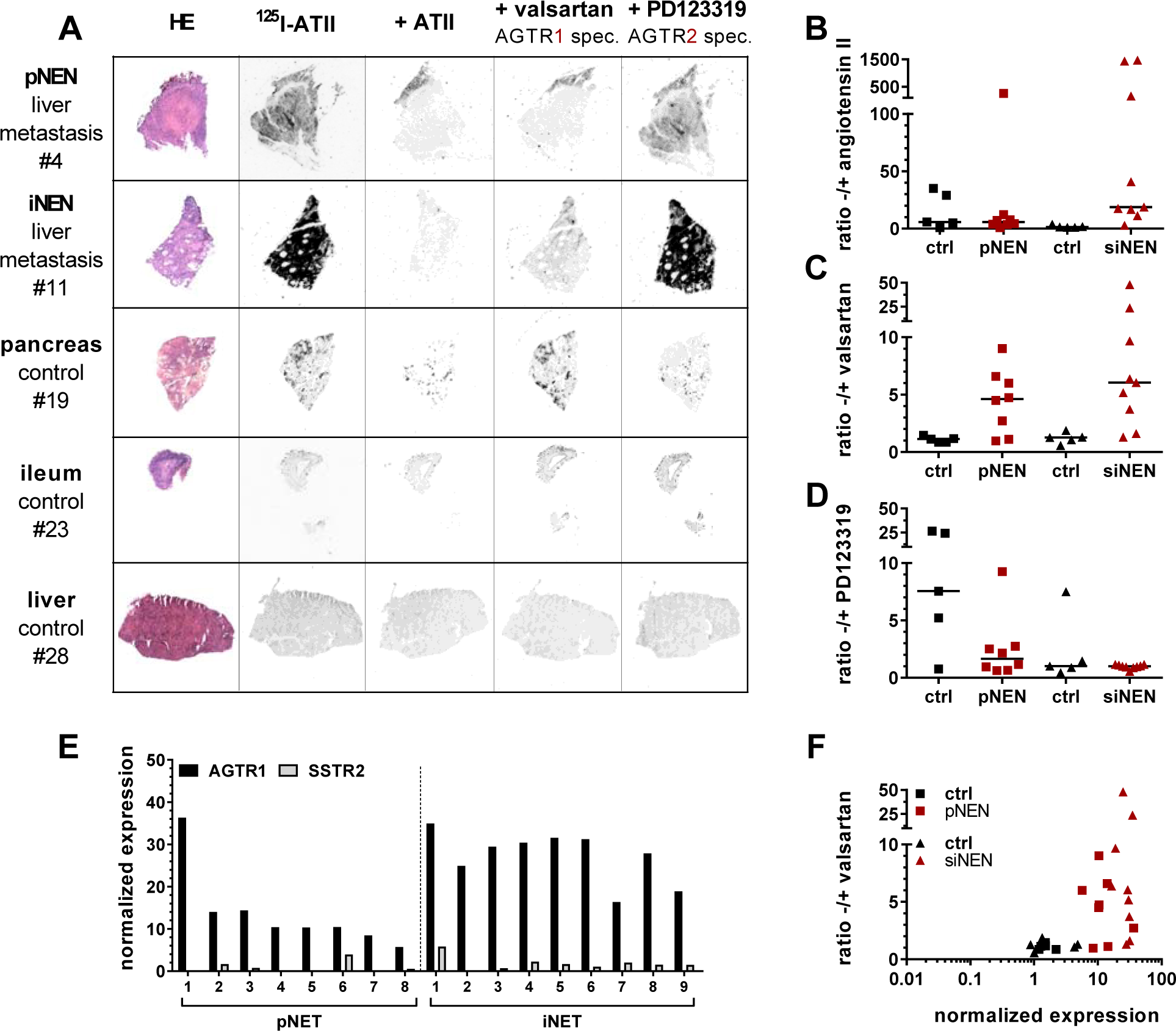
In vitro receptor autoradiography confirms increased AGTR1, not AGTR2 expression in pancreatic and small intestinal NENs. From all samples that were analyzed by RT-qPCR, 10 control and 17 NEN samples were used for autoradiographic evaluation of receptor expression. **(A)** HE staining and autoradiograms of representative pancreatic (pNEN) and small intestinal (SI-NEN) tumors as well as of pancreatic, small intestinal (SI) and liver control tissues. For each tissue, consecutive cryosections were either incubated with iodine 125 labeled angiotensin II alone (^125^I-ATII, total binding) or in the presence of additional 1 µM unlabeled ATII, AGTR1 antagonist valsartan or AGTR2 antagonist PD123319 (non-specific binding). **(B-D)** For all autoradiograms, mean signal intensities per area were calculated and are shown as ratios of total to non-specific binding for ATII (B), valsartan (C) and PD123319 (D). **(E)** Scatter plot showing the correlation of AGTR1 gene expression (x-axis, RT-qPCR) with receptor binding (y-axis, autoradiography). **(F)** For the analyzed NEN tissue samples, AGTR1 gene expression levels were compared to their SSTR2 expression, as measured with RT-qPCR (see Figure 2). Data are depicted as ratios of AGTR1 to SSTR2 expression.

Digitized autoradiograms were quantified by calculating the ratios of total binding to non-specific binding in the presence of excess ATII, valsartan or PD123319 (Fig. 3B-D). The higher a ratio, the higher the expression of the respective receptor subtype. Indeed, valsartan-related ratios were increased up to 5-fold in NEN tissues, especially in small-intestinal NENs, when compared to their respective control tissues. This confirmed a specific AGTR1 expression (Fig. 3C). To verify these findings using a different quantitative analysis, tissues were wiped off the slides after radioligand incubation and measured by a gamma counter. This alternative quantification yielded similar results (Supplementary Fig. 2), although image analysis after digitization revealed higher overall ratios. Quantitative data from autoradiography was also compared to the gene expression data for the corresponding samples. As depicted in Fig. 3E, mRNA levels could be well correlated with binding levels for most tissues (Pearson r=0.43, p≤0.01, r^2^=0.184). Whereas controls showed low mRNA and low binding levels, NENs in general had relatively high mRNA and binding levels. A similar correlation done with data from the alternative quantification confirmed this result (Supplementary Fig. 2B). Furthermore, AGTR1 mRNA levels of the samples analyzed by autoradiography were individually compared to those of SSTR2 mRNA (Fig. 3F). All these NEN tissues showed higher AGTR1 than SSTR2 mRNA levels.

After target validation in patient tissues, receptor-positive cell lines BON and H727 were further evaluated as models for AGTR1 expression and function in NEN. Confirming previous gene expression data (Fig. 2A), both cell lines exhibited significant specific binding of ^125^I-ATII that could be displaced by unlabeled ATII, whereas the other NEN cell lines QGP-1, LCC-18 and UMC-1 demonstrated only background levels of radioligand binding (Fig. 4A). Saturation radioligand binding experiments with BON and H727 cells determined K_d_ values in the (sub)nanomolar range with 0.6 nM for BON and 1.2 nM for H727 cells, respectively (Fig. 4B). In contrast, the cell lines differed in their receptor density as indicated by the B_max_ value, which was 3-fold higher for BON cells (approximately 50,000 binding sites/cell) in comparison to H727 cells (approximately 16,000 binding sites/cell). As depicted in Fig. 4C, binding of ^125^I-ATII to BON and H727 cells could be displaced in a dose-dependent fashion by ATII and the AGTR1-specific antagonist valsartan, but not by the AGTR2-specific antagonist PD123319. Calculated K_i_ values reflect the expected high affinity of ATII for AGTR1 (BON: 0.1 nM, H727: 0.2 nM). K_i_ values of valsartan were slightly higher, but still in the low nanomolar range (BON: 4.3 nM, H727: 9.4 nM). It was further shown that both cell lines retained AGTR1 expression and radioligand binding capacity during xenotransplantation and tumor growth in nude mice, as analyzed by receptor autoradiography (Fig. 4D).

**Figure 4:**
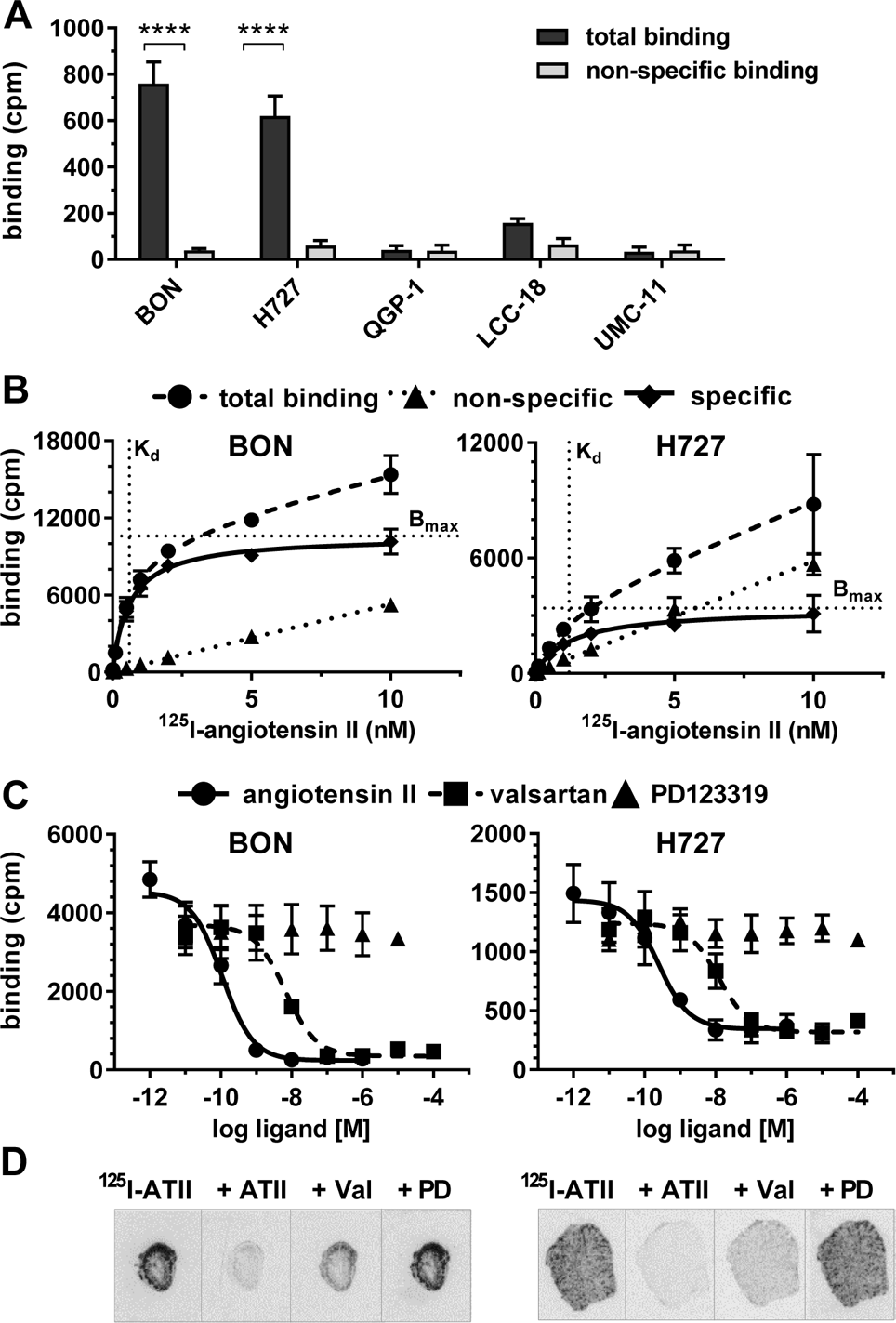
The NEN cell lines BON and H727 specifically express AGTR1, not AGTR2. **(A)** NEN cell lines were incubated with ^125^I-ATII alone (total binding) or in the presence of additional 1 µM unlabeled ATII (non-specific binding). Data represent mean ± S.E.M. (n=3). Evaluated with two-way ANOVA, ****p<0.0001. **(B, C)** Saturation (B) and competition (C) binding assays were performed for BON and H727 cells. Data represent mean ± S.E.M. (n=3). **(B)** Cells were incubated with increasing concentrations of ^125^I-ATII in absence (total binding) or presence of 1 µM unlabeled ATII (non-specific binding). Specific binding was calculated by subtraction of non-specific from total binding. **(C)** Cells were incubated with ^125^I-ATII and increasing concentrations of either unlabeled ATII, valsartan (AGTR1 antagonist) or PD123319 (AGTR2 antagonist). **(D)** Autoradiograms of BON and H727 xenografts showing total (^125^I-ATII) or non-specific binding in the presence of 1 µM ATII, valsartan or PD123319. cpm, counts per minute.

To gain an understanding of the biological function of AGTR1 in NEN cells, cellular signaling, secretion and cell viability of BON and H727 cells were evaluated upon receptor stimulation or inhibition. Functional activation of the receptor was studied by an intracellular calcium mobilization assay. While concentration-dependent response curves could be obtained for BON and H727, demonstrating functionally active receptor in these cells, QGP-1 and LCC-18 cells showed no calcium influx (Fig. 5A). Furthermore, valsartan was able to significantly diminish the induced signal, while PD123319 did not influence receptor activation, indicating selective AGTR1, not AGTR2 signaling (Fig. 5A). The AGTR1-selective antagonist valsartan inhibited ATII-mediated calcium mobilization in a concentration-dependent manner (Fig. 5B). Gα_q_ inhibition completely abolished the calcium signal, whereas β-arrestin blockage did not modulate calcium mobilization in either BON or H727 cells. Inhibition of ERK or AKT also did not change the calcium response in BON cells (Fig. 5B).

**Figure 5:**
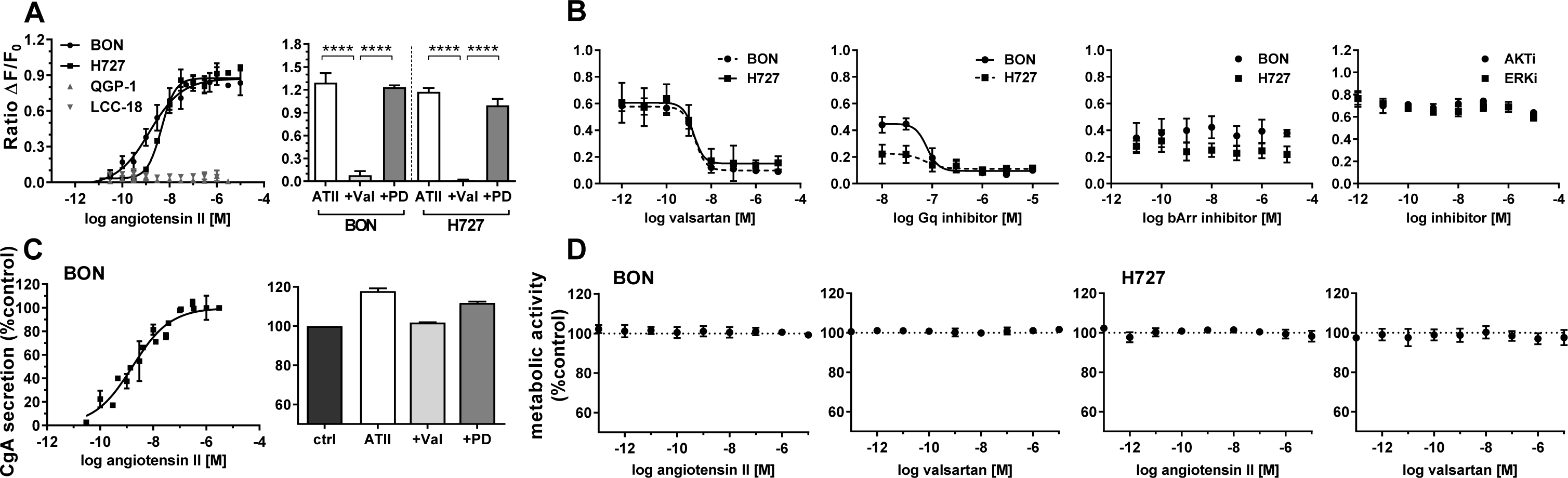
Calcium signaling, chromogranin A secretion and viability of NEN cell lines after stimulation of AGTR1. **(A)** NEN cell lines were preloaded with the calcium indicator fluo-4 and its increasing fluorescence after application of ATII was recorded (left). BON and H727 were additionally pretreated with 1 µM valsartan or PD123318 for 15 min before ATII stimulation (right). Data show mean ± S.E.M. (n≥3). Evaluated with two-way ANOVA, ****p<0.0001. **(B)** BON and H727 cells were preloaded with the calcium indicator fluo-4 and preincubated with either valsartan, G_αq_ blocker UBO-QIC, β-arrestin blocker barbadin or inhibitors for ERK or AKT (BON cells) for 15-30 min. Fluorescence intensity after application of ATII was recorded. Data show mean ± S.E.M. (n≥3) **(C)** BON cells were either treated with increasing concentrations of ATII (left) or pretreated for 15 min with 10 µM valsartan or PD123318 before application of 100 nM ATII (right). Chromogranin A (CgA) content of the supernatants was measured after 6 h (left) or 24 h (right), respectively. Data show mean ± S.E.M. (n≥2). Values were normalized on untreated controls. **(D)** BON and H727 cells were incubated with increasing concentrations of ATII or valsartan for 96 h and analyzed for their metabolic activity by addition of AlamarBlue. Data show mean ± S.E.M. (n=4).

As NENs often secrete chromogranin A (CgA), BON cell supernatants were tested for their CgA content after ATII treatment. Interestingly, CgA secretion in these cells was stimulated in a concentration-dependent manner. CgA secretion was diminished after preincubation with AGTR1 antagonist valsartan (Fig. 5C). When investigating metabolic activity as a readout for cell growth and viability, no change could be detected in BON and H727 cells 96 hours after treatment with different concentrations of agonist (ATII) or antagonist (valsartan) (Fig. 5D).

Finally, the capacity of AGTR1 to act as a target for in vivo molecular diagnostic imaging was assessed. To this end, the partial AGTR1/2 agonist saralasin was coupled via a 4,7,10-trioxatridecan-succinamic acid (TTDS) linker to the near-infrared fluorescent dye indotricarbocyanine (ITCC). Radioligand binding assays with ^125^I-ATII were conducted to determine the affinity of the fluorescent probe in vitro (Fig. 6A). In AGTR1-positive BON cells, the radioligand could be displaced by unlabeled ATII confirming its high affinity (K_i_ = 0.1 nM) as determined before (Fig. 4C). The binding curve of saralasin-ITCC was shifted to the right, indicating lower binding affinity at a K_i_ value of 246.5 nM. In contrast, receptor negative QGP-1 cells did not show binding at all. For in vivo imaging, immunodeficient nude mice were subcutaneously inoculated with BON cells on the right and QGP-1 cells on the left shoulder. Near-infrared fluorescent imaging was performed after sufficient tumor growth and the ITCC-labeled probe was tested in four different animals. Biodistribution kinetics of saralasin-ITCC of one animal after intravenous application is shown in Fig. 6B. Visual evaluation of the images indicated a selective accumulation of saralasin-ITCC in AGTR1-positive BON tumors, which proved to be statistically significant at three, four, five and six hours post-injection after quantitative analysis (Fig. 6C). Tumor-to-background ratios were 2- to 3-fold higher when compared to the image acquired prior to injection. In contrast, only low, non-significant uptake was detected in receptor negative QGP-1 tumors. Ventral images of the animals showed that the majority of the probe was excreted via the bladder. Ex vivo organ imaging 6 hours after injection confirmed the specificity of saralasin-ITCC for receptor-positive BON tumors as well as an increased accumulation in kidneys and liver (Fig. 6D). Similarly, AGTR1 small molecule antagonist valsartan was labeled with ITCC via a 1,3-diamino propane linker and used for the same type of imaging. Although the affinity of this fluorescent ligand conjugate (IC_50_=18.6 nM) was higher than for saralasin-ITCC, the accumulation of valsartan-ITCC in AGTR1-positive NEN xenograft tumors proved to be less pronounced (Supplementary Fig. 5).

**Figure 6:**
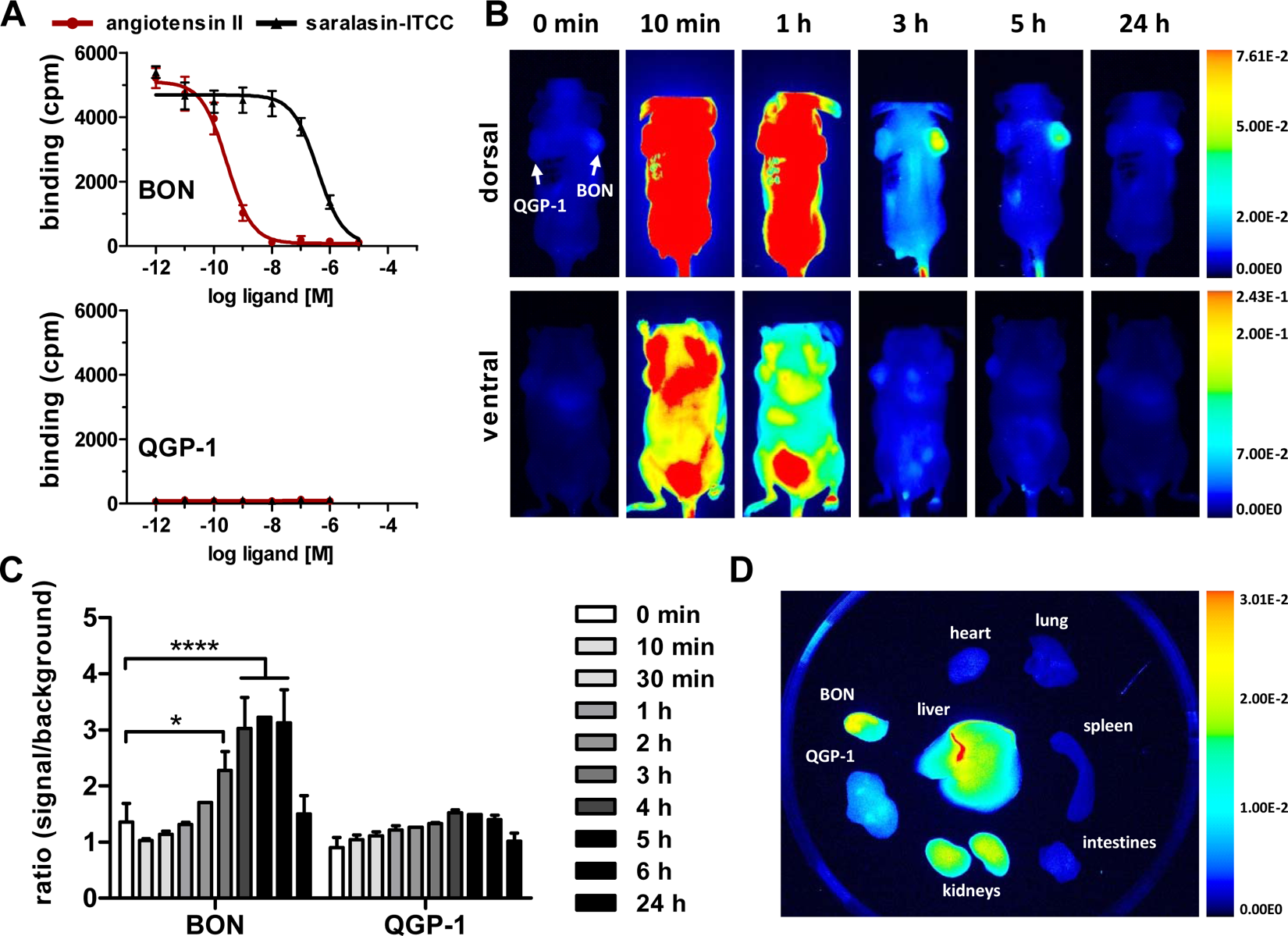
In vivo near-infrared fluorescent (NIRF) imaging with AGTR1-targeting saralasin-ITCC. **(A)** AGTR1-positive BON and AGTR1-negative QGP 1 cells were incubated with 125I-ATII and increasing concentrations of unlabeled ATII or saralasin-ITCC. Data show mean ± S.E.M. for BON (n=3-4) or mean ± S.D. for QGP 1 (n=1). **(B)** Biodistribution of 1 nmol i.v. saralasin-ITCC in a mouse model subcutaneously injected with BON (right shoulder) and QGP 1 cells (left shoulder). Images were acquired at the indicated time points before or after injection and are displayed with an equally adjusted gain for either dorsal or ventral signals. **(C)** Signals from in vivo NIRF imaging were quantified by calculating the ratio of signal (tumor) to background (neck) for saralasin-ITCC. Bars represent mean ± S.E.M. of four different animals. Evaluated with matched two-way ANOVA and Bonferroni post-hoc test, * P<0.05, **** p<0.0001. **(D)** Ex vivo NIRF imaging of tumors and organs 6 h after injection, exemplary images from one animal. cpm, counts per minute; ITCC, indotricarbocyanine.

## Discussion

ATII is the major effector of the renin-angiotensin system, regulating blood pressure and cardiovascular homeostasis. Apart from this canonical role, ATII is involved in angiogenesis, inflammation, cell proliferation, differentiation and tissue remodeling. Consequently, activation of ATII and its cognate receptors may be relevant to carcinogenesis (19, 20). So far, the ATII receptor type 1 (AGTR1) was found to be overexpressed in various cancers, such as breast (21), gastric (22) and pancreatic cancer (23, 24).

In search of alternative targets that may be used for diagnosis and treatment of NENs, this study employed cell-based screening in human NEN cells using a compound library of 998 peptides and small molecules. This approach yielded 38 confirmed hits, including ATII and a number of other GPCR ligands. These results stress the validity of activity-based screening approaches (reverse genetics) to identify new targets in oncology. In a subsequent comprehensive analysis of both AGTR1 mRNA protein expression in 123 NEN patient and 25 control samples, this study demonstrated an upregulation of AGTR1 in pancreatic and small-intestinal NENs. Tumor mRNA levels were significantly increased in comparison to healthy control tissue, especially in small-intestinal NENs. Correlation analyses revealed a positive correlation between AGTR1 and SSTR2 expression in these samples, however, a subset of patients with low SSTR2 and high AGTR1 expression could benefit from tumor targeting approaches based on AGTR1 ligands. While mRNA expression data provide important insights into differentially regulated transcript levels, the existence of sufficient receptor protein at the cell surface is a prerequisite for molecular targeting. In a previous study, AGTR1 protein expression in 44 pancreatic NENs had been evaluated by immunohistochemistry (25). This paper observed an expression in 80 % of the patients. However, the antibody employed in this paper was demonstrated to be non-specific in two studies (26, 27). In the current study, protein expression was indirectly evaluated by receptor autoradiography as an alternative more reliable method than antibody-based techniques. To discriminate between AGTR1 and AGTR2, subtype selective antagonists were co-applied for radioligand displacement and assessment of nonspecific binding. Quantification revealed an up to 5-fold higher binding of AGTR1 in pancreatic, and in particular in small-intestinal NENs. This was found to correlate well with mRNA expression for most samples.

AGTR2 expression could be detected in several pancreatic control tissues. These findings are in line with a study by Shao et al. reporting high AGTR2 levels in pancreatic islets of adult rats. (28). Sadik et al. associated the differentiation of bone-marrow derived mesenchymal stem cells into insulin producing cells with an increased AGTR2 expression (29), underlining its role in development and tissue regeneration. The distinct role of AGTR2 remains to be elucidated. Also, how AGTR1 and AGTR2 expression levels may influence tumor grade and prognosis of NENs would be of potential interest and requires further studies and integration of clinical parameters.

To identify appropriate models for investigating the impact of ATII on tumor-associated processes such as signaling, secretion and proliferation, a panel of 28 cell lines was tested for AGTR1 expression. Indeed, the two cell lines with the highest AGTR1 mRNA expression, BON and H727, were of NEN origin, whereas only one colon and one lung cancer cell line reached similarly high values. This may indicate that AGTR1 overexpression is not a general phenomenon in tumors, but may be of particular relevance to NENs. Specific AGTR1 expression in these cell lines was further confirmed by radioactive binding assays and autoradiography before applying them for in vitro and in vivo experiments.

AGTR1 is known to primarily couple to the heterotrimeric G protein G_q/11_ (20). This study confirmed for ATII-mediated mobilization of intracellular calcium in BON and H727 cells, blocked by valsartan and Gq inhibitor UBO-QIC. As Calcium acts as a versatile second messenger, physiological processes such as secretion and proliferation were investigated in more detail. Enhanced calcium channel activity had been previously associated with an increased release of the NEN marker chromogranin A (CgA) and of neurotensin (30, 31). Indeed, ATII concentration-dependently stimulated CgA secretion in BON cells. Physiological or enhanced ATII levels in the circulation of NEN patients may therefore contribute to hormone hypersecretion. Several studies showed that ATII facilitates cell proliferation in various human cancer cell lines, including breast (32), prostate (33) and pancreatic cells (34). The proposed pro-proliferative effect, however, could not be confirmed in this study, as metabolic activity was not affected in BON and H727 cells. Likewise, treatment with the AGTR1 antagonist valsartan revealed no inhibition of cell growth. One could hypothesize that BON and H727 release ATII, which may lead to autocrine signaling and enhanced cell proliferation making externally added treatment negligible. This would be in line with a previous study in which treatment with the ACE inhibitor enalapril, and thereby inhibition of ATII production, resulted in decreased BON cell growth in vitro and in vivo (25). Similarly, the effect of applied antagonists might be impaired as they have to compete with autocrine ATII binding for the receptors (35). It is also conceivable that a lack of one or more signaling components prevents the transduction of the signal from the receptor to the cell cycle machinery. Furthermore, ATII mediates its pro-proliferative signaling not only by direct stimulation of tumor cells, but also indirectly affects stromal and vascular cells (20), which are not present in most monolayer cell cultures.

Following target identification and validation, the suitability of AGTR1 as a target for optical in vivo imaging was assessed. Radiolabeled AGTR1-targeted peptides and small molecules have been primarily utilized for cardiac and cardiovascular imaging so far, to select patients for distinct treatment options and to better predict therapy response (36). To the best of our knowledge, this is the first study investigating the applicability of AGTR1-based tumor targeting. For this, the partial receptor agonist saralasin, containing three amino acid substitutions in comparison to ATII, as well as the small molecule AGTR1-antagonist valsartan were chosen. Biodistribution of the two different ITCC-labeled targeted ligands was evaluated in a xenograft mouse model. This revealed a significant accumulation of saralasin-ITCC in receptor-positive BON tumors after a few hours. Valsartan-ITCC was taken up to a lesser extent in both BON and receptor-negative QGP-1 (Supplementary Figure 5). The superior performance of saralasin-ITCC was unexpected, as the introduction of the dye molecule ITCC decreased its affinity 250-fold, compared to the published Ki value of 1 nM (37). Although a slow dissociation rate from the receptor is mostly associated with high affinity binding, both parameters do not necessarily have to correlate (38). Furthermore, saralasin exhibits a half-life of a few minutes whereas valsartan is stable for a few hours (39, 40). It is of notice though that saralasin-ITCC was quite rapidly eliminated via kidney and bladder, while valsartan-ITCC remained much longer in the mouse and was mainly excreted hepatically.

Although the tumor-to-background ratio for saralasin-ITCC was significantly increased in receptor-positive BON tumors, several modifications might further enhance probe performance. The introduction of a more hydrophobic linker between peptide and dye could result in longer retention, and hence, higher uptake and overall fluorescence intensities. Linker molecules such as aminohexanoic acid (AHX) are commonly used to increase the hydrophobicity of a molecule, whereas trioxatridecansuccinamic acid (TTDS) or polyethylene glycol (PEG) are more hydrophilic. In addition, the variation of linker length and thus the distance between peptide and near-infrared dye, might enhance the affinity of the probe (44). The proof of principle for AGTR1-based tumor imaging in this study may pave the way for a translation into diagnostic approaches like PET imaging and AGTR1-directed peptide receptor radioligand therapy (PRRT) for NENs employing chelator-based AGTR1 radiotracers.

## Acknowledgements

The authors wish to thank Ines Eichhorn and Yvonne Giesecke for expert technical assistance. This work was supported by a grant from the German Ministry of Education and Research (BMBF grants IP614 and IPT614A to CG). Samantha Exner was the recipient of a fellowship by Sonnenfeld-Stiftung.

