## Supplements for "AGTR1 is overexpressed in neuroendocrine neoplasms, regulates secretion and may serve as a target for molecular imaging and therapy"

**Supplementary Table 1: Screening library compound list**

| # | ligand |
|---|---|
| 1 | Ac2-26 (N-terminal fragment of annexin-1, lipocortin-1) |
| 2 | Ac2-12 (N-terminal fragment of annexin-1, lipocortin-1) |
| 3 | Boc-Phe-Leu-Phe-Leu-Phe |
| 4 | LL37 (Human) |
| 5 | N-Formyl-Nle-Leu-Phe-Nle-Tyr-Lys |
| 6 | N-Formyl-Met-Ile-Leu-Phe |
| 7 | N-Formyl-Met-Ile-Phe-Leu |
| 8 | N-Formyl-Met-Leu-Ile-Phe |
| 9 | N-Formyl-Met-Leu-Phe |
| 10 | N-Formyl-Met-Leu-Phe-Ile |
| 11 | N-Formyl-Met-Leu-Phe-Lys |
| 12 | N-Formyl-Met-Gln-Leu-Gly-Arg |
| 13 | WKYMVM |
| 14 | WKYMVm |
| 15 | Apelin-36 (Human) |
| 16 | Apelin-36 (Bovine) |
| 17 | Apelin-36 (Rat) |
| 18 | Apelin-19 (Human, Bovine) |
| 19 | Apelin-17 (Human, Bovine) |
| 20 | Apelin-16 (Human, Bovine) |
| 21 | Apelin-15 (Human, Bovine) |
| 22 | Apelin-13 (Human, Bovine) |
| 23 | Apelin-13, [pGlu1] (Human, Bovine) |
| 24 | Apelin (23-57) -Prepro (Human) |
| 25 | Apelin-12 (Human, Bovine) |
| 26 | Allatostatin 1 |
| 27 | Allatostatin 3 |
| 28 | Allatostatin 4 |
| 29 | Adipokinetic Hormone (AKH) (Locust) |
| 30 | Adipokinetic Hormone (Heliothis zea and Manduca sexta) |
| 31 | Adipokinetic Hormone (Taa-AKH) (Tabanus atratus) |
| 32 | Adipokinetic Hormone II (Schistocera gregaria) |
| 33 | Adipokinetic Hormone G (AKH-G) (Gryllus bimaculatus) |
| 34 | Litorin-Rohdei (AF 1 Ascaris suum) |
| 35 | Angiotensin I (Human) |
| 36 | Angiotensin I (1-9) (Human) |
| 37 | Angiotensin I, [Des-Asp1-] (Human) |
| 38 | Angiotensin II (Human) |
| 39 | Angiotensin II [Sar1] |
| 40 | D-Pro7-Angiotensin-(1-7) |
| 41 | Angiotensin II, [Sar1,Ala8] |
| 42 | Angiotensin II [Sar1 Ile8] |
| 43 | Angiotensin II (1-7) |
| 44 | Angiotensin II [Sar1Val5Ala8] (Human Canine Rat Mouse) |
| 45 | Angiotensin IV [AT-II (3-8)] |
| 46 | Angiotensin II (4-8) (Human) |
| 47 | Angiotensin II, [Des-Asp1-] (Angiotensin III) (Human) |
| 48 | Atrial Natriuretic Peptide (ANP) (Eel) |
| 49 | Atrial Natriuretic Peptide (ANP)-21 (fANP-21) (Frog) |
| 50 | Atrial Natriuretic Peptide (ANP) (1-28), Alpha (Human, Canine) |
| 51 | Atrial Natriuretic Peptide (ANP) (4-28) (Human Canine) |

52 Atrial Natriuretic Peptide (ANP) (26-55), Prepro (Human) (proANP (1-30))
53 Atrial Natriuretic Peptide (ANP) (31-67) (Prepro ANP (56-92) (Human)
54 Atrial Natriuretic Peptide (ANP) (104-123), Prepro (Human)
55 Atrial Natriuretic Peptide (ANP) (Rat)
56 Atrial Natriuretic Peptide (ANP) (4-23)-NH2, Des-[Gln18,Ser19,Gly20, Leu21,Gly22] (Rat)
57 Atrial Natriuretic Peptide (ANP) (8-33) (Rat)
58 Atriopeptin I (Rat, Rabbit, Mouse)
59 Atriopeptin II (Rat Rabbit Mouse)
60 Atriopeptin III (Rat Rabbit Mouse)
61 Atrial Natriuretic Peptide (ANP) (126-150) [Auriculin B] (Rat)
62 BNP-32 (Human)
63 BNP (1-21), Prepro (Human)
64 BNP (22-46), Prepro (Human)
65 BNP (1-46), Prepro (Human)
66 BNP-32 (Canine)
67 BNP-32 (Porcine)
68 BNP-32 (Rat)
69 BNP-45 (Rat)
70 BNP (24-45), Prepro (Rat)
71 BNP-45 (Mouse), [BNP (51-95)] (5K Cardiac Natriuretic Peptide)
72 CNP-22 (Human, Porcine, Rat)
73 CNP-53 (1-29) (Porcine, Rat)
74 DNP (Dendroaspis Natriuretic Peptide)
75 Guanylin (Human)
76 Vasopressin (AVP), [Arg8]
77 Vasopressin (AVP), [D-Arg8], desamino (DDAVP)
78 Vasopressin (AVP) (AVP) (4-9) [pGlu4 Cystine6]
79 Oxytocin
80 Oxytocin [Thr4 Gly7]
81 Bradykinin
82 Bradykinin [Sar0 D-Phe8 Des-Arg9]
83 Bradykinin [Des-Arg9]
84 Bradykinin [Lys0 D-Phe8 Des-Arg9]
85 Bradykinin [Ac-Lys0 ((-D-Nal7) Ile8 Des-Arg9]
86 Bradykinin [Des-Arg9 Leu8]
87 Bradykinin [D-Arg0 Hyp3 D-Phe7]
88 Bradykinin [D-Arg0 Hyp3 Thi5 8 D-Phe7]
89 Bradykinin [D-Arg0 Hyp2 3 Thi5 8 D-Phe7]
90 Bradykinin [D-Arg0 Hyp3 Thi5 D-Tic7 Oic8] (HOE 140)
91 Bradykinin [Lys-Lys0 Hyp3 Igl5 D-Igl7 Oic8 Des-Arg9]
92 Bradykinin [D-Arg0 Hyp3 Igl5 D-Igl7 Oic8 Des-Arg9]
93 Bradykinin [D-Arg0 Hyp3 Igl5 D-Igl7 Oic8]
94 Bradykinin [Lys0 Des-Arg9] ([Des-Arg10]-Kallidin)
95 Bradykinin [Ile-Ser-] (T-Kinin)
96 Bradykinin [Ac-Lys0-(Me-Ala6 Leu8]
97 Bradykinin [Lys0 Leu8 Des-Arg9]
98 Bombinakinin-GAP
99 Bombinakinin M
100 Bombesin Receptor AntagonistBW2258U89
101 Bombesin
102 Bombesin (6-14), [D-Phe6, Beta-Ala11, Phe13, Nle14]
103 Bombesin (6-14), [D-Tyr6, Beta-Ala11, Phe13, Nle14]
104 [D-Tyr6,(R)-Apa11,Phe13, NLe14] Bn(6-14)
105 [D-Tyr6,(S)-Apa11,Phe13, NLe14] Bn(6-14)

106 Gastrin Releasing Peptide (GRP) (Human)
107 Gastrin Releasing Peptide (GRP) (Porcine)
108 Gastrin Releasing Peptide (GRP) (14-27) (Porcine, Human)
109 Neurokinin B (Neuromedin K)
110 Neurokinin B, [D-Pro2,D-Trp6,8,Nle10] (Porcine)
111 Neurokinin B, [MePhe7]
112 Neurokinin B, [Pro7]
113 Calcitonin (salmon)
114 Calcitonin (8-32) (salmon)
115 Calcitonin II (Salmon)
116 Calcitonin III (salmon)
117 Calcitonin IV (salmon)
118 Calcitonin V (salmon)
119 Calcitonin (Human)
120 Calcitonin (Rat)
121 Calcitonin (Eel)
122 Calcitonin Receptor Stimulating Peptide (CRSP) (Porcine)
123 Calcitonin Receptor Stimulating Peptide 2 (CRSP 2) (Porcine)
124 Calcitonin Receptor Stimulating Peptide 3 (CRSP 3) (Porcine)
125 Calcitonin Receptor Stimulating Peptide, Cys0, (CRSP, Cys0) (Porcine)
126 Calcitonin Receptor Stimulating Peptide 3 (8-37) (Porcine)
127 Calcitonin Receptor Stimulating Peptide-1 (CRSP) (canine)
128 Calcitonin Gene Related Peptide (CGRP) (Rat)
129 Calcitonin Gene Related Peptide II (CGRP II) (Human)
130 Calcitonin Gene Related Peptide (CGRP) (17-37) (Human)
131 Calcitonin Gene Related Peptide (CGRP) (30-37) (Rat)
132 Calcitonin Gene Related Peptide 8-37 (Human)
133 Calcitonin Gene Related Peptide (CGRP) (8-37) (Rat)
134 Calcitonin Gene Related Peptide (CGRP) (Human)
135 Adrenomedullin (ADM), (1-52) (Human)
136 Adrenomedullin (ADM), (1-12) (Human)
137 Adrenomedullin (ADM), (13-52) (Human)
138 Adrenomedullin (ADM), (22-52) (Human)
139 Adrenomedullin (ADM), (16-52) (Human)
140 Adrenomedullin (ADM), (34-52) (Human)
141 Adrenomedullin (ADM), Pro-N-20 (PAMP-20) (Mouse)
142 Adrenomedullin (ADM), (1-50) (Rat)
143 Adrenomedullin (ADM), (24-50) (Rat)
144 Adrenomedullin (ADM), Pro-N-20 (PAMP-20) (Rat)
145 Adrenomedullin (ADM), (1-52) (Porcine)
146 Adrenomedullin (ADM), Pro-N-20 (PAMP-20) (Human)
147 Adrenomedullin (ADM), (26-52) (Porcine)
148 Adrenomedullin (ADM), (34-52) (Porcine)
149 Adrenomedullin (1-44) (Human)
150 Intermedin (Human) (hIMDL)
151 Intermedin (8-47) (Human) [hIMDS]
152 AM-2/ Intermedin (IMD) (Rat)
153 Intermedin (8-47) (IMDS) (Rat)
154 Intermedin (IMDL/AM2)(Mouse)
155 Intermedin (8-47) (IMDS) (Mouse)
156 Intermedin (17-47) (Human)
157 Intermedin (16-47), pGlu16 (Human)
158 Intermedin-53 (Human)
159 Prepro-Intermedin (25-56) (Human)

160 Prepro-Intermedin (57-92)  (IMDS) (Human)
161 Cholecystokinin (CCK) (26-33) (Non-Sulfated)
162 Cholecystokinin (CCK) (26-33)
163 Cholecystokinin (CCK) Flanking Peptide (Non-Sulfated)
164 Gastric Inhibitory Peptide (GIP) (Human)
165 Gastric Inhibitory Peptide (GIP) (Rat)
166 Progastrin, Carboxyl-Terminal Flanking Peptide of Rat (Peptide D3, FnBP)
167 Gastrin, Little (Little Gastrin) (Rat)
168 Minigastrin
169 Pentagastrin
170 Corticotropin Releasing Factor (CRF) (Human, Rat)
171 Corticotropin Releasing Factor (CRF) Antagonist (Alpha-HelicalCRF (9-41))
172 Urocortin (Human)
173 Urocortin (Rat)
174 AntiSauvagine-30 (aSvg-30)
175 Urocortin II (Mouse)
176 Stresscopin Related Peptide (SRP) (Human)
177 Stresscopin (SCP) (Human)
178 Urocortin III / SCP (3-40)-NH2] (Human)
179 Urocortin III (Mouse)
180 Urocortin II (URP) [SRP(6-43)-NH2] (Human)
181 Urocortin (D-Ser4) (Human)
182 Crustacean Cardioactive Peptide (CCAO)
183 Diuretic Hormone (DH) (Manduca sexta)
184 Endothelin-1 (ET-1) (Human,Porcine,Canine,Rat,Mouse,Bovine)
185 Endothelin-1 (ET-1) Prepro (18-50) (Human)
186 Endothelin-1 (ET-1) (22-38), Big (Human)
187 Endothelin-1 (ET-1) (6-21), [Ala1,3,11,15] (Human, Porcine, Canine, Rat, Mouse, Bovine)
188 Endothelin-1 (ET-1), Big (Human)
189 Endothelin-1 (ET-1), Big (Porcine)
190 Endothelin-1 (ET-1), Big (Rat)
191 Endothelin-3 (ET-3) (Human, Rat, Porcine, Rabbit)
192 Endothelin-3 (ET-3) (159-179), Prepro (Human)
193 Endothelin-3 (ET-3) (22-41) Amide, Big (Human)
194 BQ-610
195 BQ-788
196 Endothelin Antagonist, BQ123
197 Sarafotoxin S6a (Atractaspis engaddensis)
198 Sarafotoxin S6b (Atractaspis engaddensis)
199 Sarafotoxin S6c (Atractaspis engaddensis)
200 IRL-1620
201 Enterostatin (Human, Mouse, Rat) (APGPR)
202 Enterostatin (Porcine, Canine, Equine) (VPDPR)
203 Enterostatin (VPGPR)
204 Galanin (Bovine)
205 Galanin (Human)
206 Galanin (1-19) (Human)
207 Galanin (1-30), Prepro (Human)
208 Galanin (Porcine)
209 Galanin (65-105)-NH2, Prepro (Porcine)
210 Galanin (80-105)-NH2, Prepro (Porcine)
211 Galanin (89-105)-NH2, Prepro (Porcine)
212 Galanin (Rat)
213 Galantide

214 Galanin (1-13)-Bradykinin (2-9)
215 Galanin (1-13)-Pro-Pro-(Ala-Leu)2-Ala-Amide, M40
216 C7, Galanin (1-13)-Spantide-Amide
217 Galnon FormC-cha-Lys-MCA
218 Galanin-like Peptide (Porcine)
219 Galanin-like Peptide (Human)
220 Galanin-like Peptide (Rat)
221 Galanin-like Peptide (36-60) (Human)
222 Growth Hormone Releasing Factor (GHRF) (Human)
223 Growth Hormone Releasing Factor (GHRF) (Mouse)
224 Growth Hormone Releasing Factor (GHRF) (1-29) Amide (Human)
225 Growth Hormone Releasing Factor (GHRF) (Rat)
226 Growth Hormone Releasing Factor (GHRF) (1-29) Amide, [Ac-Tyr1,D-Phe2]
227 Growth Hormone Releasing Factor (GHRF) (1-29) Amide, [N-Ac-Tyr1,D-Arg2] (Human)
228 Growth Hormone Releasing Factor (GHRF) (1-32) Amide, [His1,Ile27] (Human)
229 Growth Hormone Releasing Factor (GHRF) (30-44) Amide (Human)
230 JV-1-36 (Human)
231 JV-1--38 (Human)
232 Growth Hormone Releasing Factor-6 (GHRF-6)
233 Growth Hormone Releasing Peptide-6 (GHRP-6) [D-Lys3]
234 Ghrelin (Human)
235 Ghrelin (Rat Mouse)
236 Ghrelin [Des-Octonyl-Ser3] (Human)
237 Ghrelin [Des-Octanoyl3] (Rat Mouse)
238 Ghrelin (86-117) -Prepro (Human)
239 Ghrelin (52-85) -Prepro (Human)
240 Ghrelin (86-117) -Prepro (Rat Mouse)
241 Ghrelin (52-85) -Prepro (Rat Mouse)
242 Ghrelin (3-28) (Rat Mouse)
243 Ghrelin (1-14) (Human)
244 Ghrelin (1-14) (Human) (Des-Octanoyl3)
245 Ghrelin (1-5)-NH2
246 Ghrelin (1-5)-NH2 (Des-Octanoyl3)
247 Ghrelin C-terminal Hexapeptide
248 Ghrelin (17-28) (Human Rat)
249 Ghrelin [Des-Q14-] (Rat Mouse)
250 Ghrelin [Des-Gln14] (Des-Octanoyl3) (Rat Mouse)
251 Ghrelin [Des-Octanoyl] (1-18) Motilin-Related Peptide (Human)
252 Ghrelin [Des-Octanoyl] (1-18) Motilin-Related Peptide (Rat Mouse)
253 Ghrelin (1-11) (Rat Porcine Mouse)
254 Ghrelin (Canine)
255 Ghrelin [Des-Gln14] (Canine)
256 Ghrelin (Porcine)
257 Ghrelin [Dap-Octanoyl 3] (Human)
258 Ghrelin-21 (Eel) (O-n-Octanoyl-Ser3)
259 Ghrelin-21-C10 (Eel) (O-n-Decanoyl-Ser3)
260 Ghrelin (Tilapia)
261 Ghrelin, Gln28 (Human) (O-n-Octanoyl-Ser3)
262 Ghrelin (24-58), Gln51, Prepro (Human)
263 Ghrelin (1-9), Gly8 (O-n-Octanoyl-Ser3)
264 Ghrelin (1-7)-Lys Amide (O-n-Octanoyl-Ser3)
265 Ghrelin (1-4) Amide (O-n-Octanoyl-Ser3)
266 Ghrelin (1-18) Amide (Human) (O-n-Octanoyl-Ser3)
267 Ghrelin Amide (Tilapia) (O-n-Decanoyl-Ser3)

268 Ghrelin (1-9) Amide (Human) (O-n-Octanoyl-Ser3)
269 Ghrelin (1-14) Amide (Human) (O-n-Octanoyl-Ser3)
270 Ghrelin (1-7) Amide (Human) (O-n-Octanoyl-Ser3)
271 Ghrelin (1-11) Amide (Human) (O-n-Octanoyl-Ser3)
272 Ghrelin (1-18) Free Acid (Human) (O-n-Octanoyl-Ser3)
273 Ghrelin (52-75), Prepro, (Human)
274 Ghrelin (1-27), (O-n-Octanoyl-Ser3) (Human)
275 Ghrelin (1-27), (O-n-Decanoyl-Ser3) (Human)
276 Ghrelin, (O-n-Decanoyl-Ser3) (Human)
277 Ac-Glu-12-Ile
278 Glucagon (Human)
279 Glucagon (78-107)-NH2 Prepro (Glucagon-Like Peptide-1 (GLP-1) (7-36)) (Human)
280 Glucagon-Like Peptide-1 (GLP-1) (7-37) (Human)
281 Glucagon (72-108)-NH2 Prepro (Glucagon-Like Peptide-1 (GLP-1)) (Human)
282 Glucagon (126-159) Prepro (Glucagon-Like Peptide-2 (GLP-2)) (Human)
283 Exendin-3 (Heloderma horridum)
284 Exendin-3 (3-39) (Heloderma horridum)
285 Exendin-4 (Heloderma suspectum)
286 Exendin-4 (3-39)
287 Glucagon (Dogfish, Scyliorhinus canicula)
288 Glucagon 37 (Oxyntomodulin)
289 Glucagon (19-29) (Human)
290 GLP-2 (Glucagon-Like Peptide-2) [Des-Arg34]
291 Melanin Concentrating Hormone (MCH) [Phe13,Tyr19] (Human, Mouse, Rat)
292 Melanin Gene Overprinted Polypetide-14 (MGOP-14)
293 Melanin Gene Overprinted Polypetide-27 (MGOP-27) (Human)
294 Melanin Concentrating Hormone (MCH) (Human, Mouse,Rat)
295 Melanin Concentrating Hormone (MCH) (Salmon)
296 NEI-MCH [Prepro-MCH (131-165)]
297 Neuropeptide GE (Human)
298 Neuropeptide GE (Rat)
299 Neuropeptide GE (Mouse)
300 Neuropeptide EI (NEI)
301 Motilin (Human, Porcine)
302 Motilin (Canine)
303 Alpha-Melanocyte Stimulating Hormone see MSH, Alpha
304 MSH, [Des-Acetyl]-Alpha
305 MIF
306 MSH, Beta (Human)
307 MSH, Beta (Monkey)
308 MSH, Gamma
309 MSH, Gamma1
310 MSH, Lys-Gamma1
311 MSH, Gamma3
312 MSH, Alpha, [Nle4, D-Phe7]
313 MSH-BETA (Porcine)
314 MT II
315 SHU 9119
316 HS 014
317 JKC363
318 HS 024
319 JKC366 (Cyclic-[Mpr3-Nal4-Arg5-D-Nal7-Cys11-NH2]-alpha-MSH (3-11))
320 MT II (D-Tyr4)
321 MC-4R agonist

322 JKC-372 Cyclic-[Mpr11,D-Phe14,Cys18,Asp22-NH2]-β-MSH (11-22
323 Agouti (1-40)-Amide (Human)
324 Agouti-Related Protein (AGRP) (25-51) (Human)
325 Agouti-Related Protein (AGRP) (54-82) (Human)
326 Agouti-Related Protein (AGRP) (83-132)-NH2 (Human)
327 Agouti-Related Protein (AGRP) Form C-NH2 (Human)
328 Agouti-Related Protein (AGRP) (82-131)-NH2 (Mouse)
329 Agouti Protein (93-132)-NH2 (Mouse)
330 Neuromedin U-25 (Human)
331 Neuromedin U (Rat)
332 Neuromedin U-8 (Porcine)
333 Neuromedin U-25 (Porcine)
334 Neuromedin U-9 (Guinea Pig)
335 Neuromedin C (Porcine)
336 Neuropeptide Y (NPY) (Human, Rat)
337 Neuropeptide Y (NPY) Free Acid (Human)
338 Neuropeptide Y (NPY) (Porcine)
339 Neuropeptide Y (NPY) [Leu31,Pro34] (Porcine)
340 Neuropeptide Y (NPY) (2-36) (Porcine)
341 Neuropeptide Y (NPY) (13-36) (Porcine)
342 Neuropeptide Y (NPY) (16-36) (Porcine)
343 Neuropeptide Y (NPY) (18-36) (Porcine)
344 Neuropeptide Y (NPY) (20-36)
345 Neuropeptide Y (NPY) [D-Trp34] (Human Rat)
346 Neuropeptide Y (NPY) [D-Trp32] (Porcine)
347 Neuropeptide Y (NPY) (3-36) (Porcine)
348 Peptide YY (PYY) (Human)
349 Peptide YY (PYY) (Porcine, Rat)
350 Peptide YY (PYY) (3-36) (Human)
351 Peptide YY (PYY) (3-36) (Porcine, Rat)
352 Peptide YY (PYY) (13-36) (Porcine, Rat)
353 Pancreatic Polypeptide (Human)
354 Pancreatic Polypeptide (Rat)
355 Pancreastatin (Chromograninin A (250-301)-Amide) (Human)
356 Pancreastatin (24-52) (hPST-29) (Human)
357 Pancreastatin (Porcine)
358 Pancreastatin (Chromograninin A (264-314)-Amide) (Rat)
359 Parastatin (1-19) (Chromograinin A (347-365)) (Porcine)
360 Neurotensin
361 Kinetensin (Neurotensin Related Peptide)
362 Neurotensin [D-Trp11]
363 Neurotensin (1-7)
364 Neurotensin (8-13)
365 Neurotensin (8-13), Acetyl
366 Neurotensin (8-13), [Lys8,Asn9] (LANT-6)
367 Neuromedin N (Porcine)
368 Orexin A (Human, Rat, Mouse)
369 Orexin B (Human)
370 Orexin B (Mouse, Rat)
371 Orexin B (3-28) (Mouse, Rat)
372 Orexin B - [Ala11 - D-Leu15](Human)
373 Orexin A (16-33)-NH2 (Human, Mouse, Rat)
374 Orexin A (Human, Rat, Mouse), [Cys(Acm)6,12]
375 Hypocretin-1 (Mouse)

376 Hypocretin-2 (Mouse)
377 Orexin B (6-28) (Human)
378 Orphanin FQ/Nociceptin (Ox, Mouse, Rat, Human)
379 Orphanin FQ/Nociceptin (1-7)
380 Orphanin FQ/Nociceptin (85-119) (Prepro) [Nocistatin-35 (Rat)]
381 Orphanin FQ/Nociceptin (141-157) -Prepro
382 Orphanin FQ/Nociceptin [Tyr14]
383 Orphanin FG/Nociceptin [Des-Phe1]
384 Orphanin FQ/Nociceptin (1-11)
385 Orphanin FQ/Nociceptin (85-119) [Tyr0] -Prepro (Nocistatin-35 [Tyr0] (Rat))
386 Orphanin FQ/Nociceptin [Phe=Gly) -(1-13)-NH2
387 Orphanin FQ/Nociceptin (1-13)-NH2
388 Orphanin FQ/Nociceptin (173-181) -Pro (Rat)
389 Orphanin FQ (154-181) -Pro (Rat) [Orphanin FQ (Murine) (160-187) -Prepro]
390 Orphanin FQ (111-127) -Prepro [Nocistatin/PNP-3]
391 PNP-3 [Tyr0] (Nocistatin [Tyr0]) (Bovine)
392 Nocistatin [Lys14-E-(Tyr)]/PNP-3 [Lys14-E-(Tyr)]
393 PNP-3-8P (Bovine)
394 PNP-2/3 (Mouse) (Nocistatin-41 (Mouse))
395 Nociceptin Antagonist [AC-Arg-Tyr-Tyr-Arg-Ile-Lys-Amide]
396 Orphanin FQ/Nociceptin [N-Phe1] (1-13)-Amide
397 Dynorphin A (1-8) (Porcine)
398 Dynorphin A (1-10) (Porcine)
399 Dynorphin A (1-10)-Amide (Porcine)
400 Dynorphin A (1-13) (Porcine)
401 Dynorphin A Amide (Porcine)
402 Dynorphin A (1-6) (Porcine) (Leu-Enkephalin-Arg) (alpha-Neo-Endorphin (1-6))
403 Dynorphin A (1-7) (Porcine) (Leu-Enkephalin-Arg-Arg)
404 Dynorphin A (1-13) Amide (Porcine)
405 Dynorphin A (3-13) (Porcine)
406 Dynorphin A (6-17) (Porcine)
407 Dynorphin A (9-17) (Porcine)
408 Dynorphin B/Rimorphin (Porcine)
409 Endorphin, Beta-Neo
410 Endorphin, Ac-Alpha (Lipotropin (61-76), Ac-Alpha)
411 Alpha Endorphin (Beta-Lipotropin (61-76)
412 Xen-Dorphin-1B
413 Xen-Dorphin-1A
414 Endorphin Beta (Camel Bovine Ovine)
415 Endorphin Ac-Beta (Human)
416 Endorphin, Beta (Human)
417 Endorphin Beta (2-31) (Human)
418 Endorphin [Des-Tyr1]-Beta (Human) (Beta-Lipotropin 62-91)
419 Endorphin (1-26), Beta (Human)
420 Endorphin (1-27), Beta (Human)
421 Endorphin (18-31), Beta (Human)
422 Endorphin (6-31) Beta (Human)
423 Endorphin, Beta (Rat)
424 Endorphin, Gamma (Beta-Lipotropin (61-77))
425 Lipotropin (61-64) Beta
426 Lipotropin (61-69) Beta
427 Lipotropin (88-91) Beta
428 Receptorphin
429 Enkephalin A (104-129)-NH2 Pro (Amidorphin) (Bovine)

430 BAM (8-22)
431 BAM-12P (Bovine)
432 BAM -18P
433 BAM-22P (Bovine)
434 Somatostatin Analog (CTOP)
435 CTAP
436 Enkephalin [D-Pen2 Pen5]
437 Leu-Enkephalin
438 Delta-Receptor Peptide (DSLET) (Leu-Enkephalin-Thr, [D-Ser2])
439 Leu-Enkephalin-Thr, [D-Thr2] (DTLET)
440 Met-Enkephalin
441 Spinorphin (Bovine)
442 Enkephalin [D-Pen2 pCl-Phe4 D-Pen5] (pCL-DPDPE)
443 Met-Enkephalin-Arg-Gly-Leu (Tyr-Gly-Gly-Phe-Met-Arg-Gly-Leu)
444 Met-Enkephalin-Arg-Phe (Tyr-Gly-Gly-Phe-Met-Arg-Phe)
445 Metorphinamide (Adrenorphin)
446 3200-dalton Adrenal Peptide E (Bovine)
447 Peptide F (Bovine)
448 Tyr-Gly-Gly-Phe-Met-Arg-Phe-NH2
449 Spinorphin (Bovine)
450 Enkephalin [Des-Tyr1 D-Pen2 Pen5]
451 Enkephalin [D-Pen2 5]
452 Casomorphin, Beta
453 Casomorphin (1-4) Amide, Beta (Morphiceptin)
454 Endomorphin-1, (Tyr-Pro-Trp-Phe-NH2)
455 Endomorphin-2, (Tyr-Pro-Phe-Phe-NH2)
456 Deltorphin I, [D-Ala2]
457 Hemorphin-4 (H-4) (Bovine, Human)
458 Casoxin D
459 PACAP27-NH2 (Human, Ovine, Rat)
460 PACAP (6-27) (Human, Ovine, Rat)
461 PACAP38 (Human, Ovine, Rat)
462 PACAP (6-38) (Human, Ovine, Rat)
463 PACAP-Related Peptide (PRP) (12-29) (Rat)
464 PACAP38 (31-38) (Human, Ovine, Rat)
465 PACAP-Related Peptide (PRP) (Human)
466 Vasoactive Intestinal Peptide (VIP) (Human, Porcine, Rat)
467 Vasoactive Intestinal Peptide (VIP) (1-12) (Human, Porcine, Rat)
468 Vasoactive Intestinal Peptide (VIP) (10-28) (Human, Porcine, Rat)
469 Vasoactive Intestinal Peptide (VIP) Antagonist
470 Vasoactive Intestinal Peptide (VIP) Receptor Antagonist
471 VIP Receptor Binding Inhibitor, L-8-K
472 VIP1 Agonist: [Lys15, Arg16, Leu27]-VIP (1-7)-GRF (8-27)
473 VIP1 Antagonist: [Ac-His1, D-Phe2, Lys15, Arg16, Leu27]-VIP (1-7)-GRF (8-27)
474 VIP1 Agonist Secretin [Arg16] (Chicken)
475 VPAC2 Ligand
476 PHI-27 (PHI) (Porcine)
477 PHI (Rat)
478 PHI (1-27)-Gly (Rat)
479 PHM-27 (PHI) (Human)
480 PHM-VIP Space Peptide (PHM (111-122)/Prepro VIP)
481 PHM (156-170) (Prepro VIP)
482 Helospectin, I
483 Prolactin-Releasing Peptide-31 (PrRP-31) (Human)

484 Prolactin-Releasing Peptide-20 (PrRP-20) (Human)
485 Prolactin-Releasing Peptide-31 (PrRP-31) (Rat)
486 Prolactin-Releasing Peptide-20 (PrRP-20) (Rat)
487 Prolactin-Releasing Peptide-31 (PrRP-31) (Bovine)
488 Prolactin-Releasing Peptide-20 (PrRP-20) (Bovine)
489 PSFHSWS-Amide (Aleucokinin-like Peptide Receptor ligand)
490 Parathyroid Hormone (PTH) (1-44) (Human)
491 Parathyroid Hormone (PTH) (1-38) (Human)
492 Parathyroid Hormone (PTH) (1-34) [Nle8,18,Tyr34] (Human)
493 Parathyroid Hormone (PTH) (1-34) (Human)
494 Parathyroid Hormone (PTH) (1-34)-Amide, [Nle8,18,Tyr34] (Human)
495 Parathyroid Hormone (PTH) (3-34)-Amide, [Nle8,18,Tyr34] (Human)
496 Parathyroid Hormone (PTH) (39-68) (Human)
497 Parathyroid Hormone (PTH) (44-68) (Human)
498 Parathyroid Hormone-Related Protein (PTH-RP) (1-34) (Human)
499 Parathyroid Hormone-Related Protein (PTH-RP) (1-36)-Amide, [Tyr36] (Chicken)
500 Parathyroid Hormone-Related Protein (PTH-RP) (1-34) Amide, [Tyr34] (Human)
501 Parathyroid Hormone-Related Protein (PTH-RP) (1-37) (Human)
502 Parathyroid Hormone-Related Protein (PTH-RP) (7-34) Amide (Human)
503 Parathyroid Hormone-Related Protein (PTH-RP) (7-34) Amide, [Asn10,Leu11] (Human)
504 Parathyroid Hormone-Related Protein (PTH-RP) (7-34) Amide, [Leu11,D-Trp12] (Human)
505 Parathyroid Hormone-Related Protein (PTH-RP) (38-64) Amide (Human)
506 Parathyroid Hormone-Related Protein (PTH-RP) (67-86) Amide (Human)
507 Parathyroid Hormone-Related Protein (PTH-RP) (107-111) (Human)
508 Parathyroid Hormone-like Peptide (PLP) (140-173) (Human)
509 TIP 39
510 TIP (24-39) (Bovine)
511 TIP (7-39) (Bovine)
512 TIP39 (Mouse)
513 RFRP-3 (Human)
514 RFRP [Prepro] (56-92)-Amide (Human)
515 RFRP-1 (Rat, Mouse)
516 RFRP [Prepro] (115-131)-amide (Human)
517 RFRP-1 (Human)
518 RFRP-2 (Human)
519 RFRP [Prepro] (103-125)-amide (Mouse)
520 RFRP [Prepro] (115-131)-Amide (Human)
521 Metastin (Human)
522 Metastin (27-54)-NH2+D1220 (Human)
523 Metastin (45-54)-NH2 (Human)
524 pGlu26-Metastin (26-54)-NH2 (Human)
525 KISS-1 (125-145)
526 KiSS-1 (110-119)-NH2 (Mouse)
527 KiSS-1 (68-119)-NH2 (Mouse)
528 Antho-Rwamide II
529 Antho-RWamide I
530 Lys4-Antho-RWamide I
531 Trp7-NF1
532 Gly4,Trp7-NF1
533 Asn3-Antho-RWamide II
534 Glu1,Lys4-Antho-RWamide I
535 Ser3-Antho-RWamide II
536 Glu1, Ala4-Antho-RWamide I
537 Tyr3-Antho-RWamide II

538 Antho-RWamide I [Ala4]
539 P518/Hypothetical Protein XP_294524 (171-196)-NH2/QRFP-26 (Human)
540 P550/Hypothetical Protein XP_149121 (97-122)-NH2 (Mouse)
541 Hypothetical Protein XP_294524 (154-196)-NH2 / QRFP-43 (Human)
542 QRFP-43 (Rat)
543 QRFP-43 (Mouse)
544 QRFP-26 (13-26)(Rat)
545 QRFP-7 (Human)
546 Neuropeptide AF (huNPAF) (Human)
547 Neuropeptide FF (huNPFF, NPSF) (Human)
548 Neuropeptide AF (Bovine) (bNPAF, A18Fa)
549 Neuropeptide FF (Bovine) (bNPAF, F8Fa)
550 Morphine Modulating Peptide, C-Terminal Fragment
551 FMRF-Like Peptide
552 F1 Peptide (Lobster)
553 Tyr-Met-Arg-Phe-NH2
554 Secretin (Human)
555 Secretin (Mouse)
556 Secretin (Rat)
557 Secretin (Canine)
558 Somatostatin
559 Somatostatin (25-34) Prepro- (Antrin, Pro-Somatostatin (1-10))
560 Somatostatin (1-32) Pro (Somatostatin, N-Terminal Prepro) (Porcine)
561 Somatostatin-25
562 Somatostatin-28
563 Somatostatin-28 (1-12)
564 Somatostatin-28 (1-14)
565 Somatostatin Analog, RC-160
566 Somatostatin Analog (CTOP)
567 Octreotide
568 Cortistatin-29 (Rat)
569 Cortistatin-29 (1-13) (Rat)
570 Cortistatin-17 (Human)
571 Cortistatin-29 (Human)
572 Substance P
573 Substance P, [Tyr8]
574 Substance P, [D-Arg1,D-Phe5,D-Trp7,9,Leu11]
575 Substance P (6-11), Ac-[Arg6,Sar9,Met(O2)11]
576 Senktide, Selective Neurokinin B Receptor Peptide
577 Neurokinin A, (Neuromedin L), (Substance K)
578 Neurokinin A (4-10), [Tyr5,D-Trp6,8,9,Arg10] (MEN 10, 207)
579 Endokinin A/B
580 Endokinin C
581 Endokinin D
582 gamma Tachykinin 4 (g-TAC4) (30-61)-NH2 (Human)
583 gamma Tachykinin 4 (g-TAC4)(32-50)-NH2 (Human)
584 Hemokinin-1 (HK-1) (Mouse, Rat)
585 HK-1 (4-11) (Human)
586 Hemokinin-1 (HK-1) (Human)
587 Thrombin Receptor Agonist
588 Thrombin Receptor Agonist Peptide (TRAP1-6)
589 Thyrotropin Releasing Hormone (TRH)-Gly-Lys-Arg
590 Prepro-TRH 53-74
591 Prepro-TRH (83-106)

592 Prepro-TRH (115-151)
593 Prepro-TRH (160-169)
594 Prepro-TRH (178-199)
595 Vasotocin, [Arg8]
596 Urotensin II (Human)
597 Urotensin II (Frog)
598 Urotensin II (Mouse)
599 Urotensin II (Rat)
600 Urotensin II (87-110) -Prepro (Human)
601 SB-710411 (Urotensin II Receptor Antagonist)
602 URP / Urotensin II-Related Peptide (Human, Mouse, Rat)
603 Urotensin II (3-11) (Human)
604 ACTH (Human)
605 ACTH (Rat)
606 ACTH (1-24) (Human)
607 ACTH (18-39) [Corticotropin Like Intermediate Lobe Peptide (CLIP)] (Human)
608 ACTH [D-Arg8-] (4-10)
609 ACTH (7-38) (Corticotropin Inhibiting Peptide, CIP) (Human)
610 Leptin (Human)
611 Leptin (Mouse)
612 Leptin (57-92) (Human)
613 Leptin (22-56) (Human)
614 NEP1-40
615 Neuropeptide F (Monieza expansa)
616 Proctolin
617 Icilin
618 HezDH
619 Neuropeptide B-23 (NPB-23), Des-Br, (Human)
620 Neuropeptide B-29 (NPB-29), 6-Br, (Human)
621 Neuropeptide W-23 (Human)
622 (Des-Br)-NPB-29 (Human)
623 [6-Br-L-Trp]-NPB-29 (Mouse)
624 NPB-29, Des-Br, (Mouse)
625 [6-Br-D-Trp]-NPB-29 (Rat)
626 NPB-29 (Rat)
627 NPB-29, Des-Br, (Rat)
628 NPW-30 (Human)
629 NPW-30 (Rat)
630 NPW-23 (Mouse, Rat)
631 NPW-23 (Porcine)
632 [6-Br-D-Trp1]-NPB-29 (Human)
633 NPB-29 (Bovine)
634 NPB-29, Des-Br, (Bovine)
635 D-Trp1-NPB-29 (Bovine)
636 [6-Br-D-Trp1]-NPB-29 (Mouse)
637 [6-Br-D-Trp1]-NPB-29 (Bovine)
638 Insulin (Human), Recombinant
639 C-peptide (Human)
640 C-Peptide (Rat)
641 INSL3 (Human)
642 INSL4 (Human)
643 INSL6 (Human)
644 INSL7/ H3 Relaxin (Human)
645 INSL 3 (Mouse)

646 [Cys(Acm)10]-INSL-3 B-Chain (Mouse)
647 INSL 5 (Mouse)
648 INSL-6 C-Peptide (153-168) (Human)
649 INSL-6 Prepro-C-Peptide (121-148) (Human)
650 INSL 3 C Peptide / INSL 3 Prepro (58-105) / Insulin-like 3 C Peptide / Insulin-like 3 Prepro (58-105) (Human
651 INSL 4 A Chain / Insulin-like 4 A Chain (Human)
652 Prokineticin 2 (PK2) (82-108), Cys(Acm)86, 94 (Human)
653 Prokineticin 2 (PK2) (82-108), Cys86-94, 88-104 (Human)
654 Prokineticin 2 (PK2) (89-108), Cys(Acm)94 (Human)
655 Prokineticin 1 (PK1) (81-105), Cys (Acm)96 (Human)
656 Prokineticin 1 (PK1) (75-105), Cys (Acm)78, 96 (Human)
657 Growth Factor Antagonist Platelet-Derived
658 Gln-Alltostatin C (Gln-AlC)
659 pGlu-Alltostatin C (pGlu-AlC)
660 AF9
661 Pheromone Biosynthesis-Activating Neuropeptide (PBAN Hez)(Heliothis zea)
662 PBAN-1 (Bombyx mon)
663 Leucopyrokinin (LPK)
664 Myomodulin
665 Growth-blocking peptide
666 Xenopsin 25 (Xinin 25)(Human)
667 Secretoneurin (Rat)
668 Chemerin/TIG2 (145-157), Tyr145, Phe149, (Human)
669 TIG-2 (135-154) (Human)
670 [Cys(Acm)98,117]-TIG-2 (98-137) (Human)
671 TIG-2/ Chemerin (145-157) (Human)
672 Sun A, C-terminal 24 ammo acid polypeptide
673 Sun B, C-terminal 20 ammo acid polypeptide
674 Neuropeptide S (Mouse)
675 Neuropeptide S (Rat)
676 Neuropeptide S (Human)
677 Neuropeptide S (1-10) (Rat)
678 Neuropeptide S (1-15) (Mouse)
679 Neuropeptide S (1-15 ) (Rat)
680 Neuropeptide S (1-12) (Human)
681 Neuropeptide S (4-20) (Human)
682 WRWWWW
683 F2L/HBP (1-21) Acetylated (Human, Porcine, Canine)
684 F2L/HBP (1-21) Nonacetylated (Human, Porcine, Canine)
685 F2L/HBP (1-21) Nonacetylated (Rat, Mouse)
686 u PAR (84-95)
687 u PAR (84-95), scrambled peptide
688 SHAAG Peptide
689 rattin
690 Humanin HN (M)
691 Humanin HN (N)
692 Humanin (N) (Rat)
693 Humanin, Pro8, HN (N)
694 Humanin, Gly14, HN (N)
695 Neuromedin S (17-33) Amide (Human)
696 Neuromedin S (20-36) Amide (Mouse, Rat)
697 Neuromedin S (Human)
698 Neuromedin S (Rat)
699 Neuromedin S (Mouse)

700 Prepro-Neuromedin S (70-103) (Human)
701 Prepro-Neuromedin S (70-103) (Rat)
702 Prepro-Neuromedin S (70-103) (Mouse)
703 Prepro-Neuromedin U (104-136) (Human)
704 Beta-Alanine
705 Adenine
706 GnRH-II
707 Head Activator Neuropeptide
708 Obestatin (Human, Monkey)
709 Obestatin-Gly-Lys (Human, Monkey)
710 Obestatin (Mouse, Rat)
711 Adiponectin (17-41) (Human)
712 Adiponectin (17-41) (Rat)
713 Beta-Defensin 2
714 Beta-Defensin 8 (Mouse)
715 Drm-AST-3
716 Tunicate-3
717 Tunicate-5
718 Tunicate-6
719 N-Formyl-MDGCEL
720 pGlu-Gln-Pro-Amide
721 Big Dynorphin [Prodynorphin (209-240)](Porcine)
722 Agouti-Related Protein (25-82) / AGRP (25-82) (Human)
723 Endothelin-1, Prepro (169-212) (Human) /PSW44
724 Prepro-Endothelin-1 (145-162)-Amide / Prepro-ET-1 (145-162)-Amide (Human)
725 Prepro-Endothelin-1 like Peptide (109-130) / Prepro-ET-1 like Peptide (109-130) (Human)
726 Prepro-Endothelin-1 (131-142) / Prepro-ET-1 (131-142) (Human)
727 Prepro-Endothelin-1 (109-125) Amide / Prepro-ET-1 (109-125) Amide (Human)
728 Prepro-Endothelin-1 (93-108) / Prepro-ET-1 (93-108) (Human)
729 [Cys0]-Prepro-Endothelin-1 (168-181) / [Cys0]-Prepro-ET-1 (168-181) (Human)
730 Sialorphin (Rat)
731 Opiorphin (Human)
732 Enkephalin (219-229) -Pro
733 Enkephalin Octapeptide - Pro
734 Enkephalin (239-262) -Pro
735 Enkephalin (239-262) -Pro
736 Virp5 (4-11)(Rat) / Anti-Verotoxin Antibody ImmunoReactive Peptide
737 Virp5 (19-26) / P-19 (Rat)
738 Virp5 (36-43) / P-36 (Rat)
739 Brevinin-1
740 Relaxin-2 (Human)
741 [Des-C-Peptide]-IGF 1 (Human)
742 Insulin-1 (Mouse)
743 MGF / C-Terminal Peptide of IGF-1 Ec (Human)
744 Insulin-1 (Rat)
745 Prepro-Urotensin II (87-104) (Rat)
746 Prepro-Urotensin II (71-84) (Human)
747 Prepro-Urotensin II (41-68) (Human)
748 Prepro-Urotensin II (21-40) (Human)
749 Prepro-Urotensin II (87-110) (Human)
750 Urotensin II (4-11) (Human)
751 Urotensin II (5-11) (Human)
752 UFP-803 / [Pen5, D-Trp7, Dab8]-Urotensin II (4-11) Cyclic
753 [Ala13]-Apelin-13 (Human, Bovine)

754 [pGlu1,Ala13]-Apelin-13 (Human, Bovine)
755 Apelin-28 (Human)
756 Prepro-Apelin (33-46) (Human)
757 Neuropeptide S (1-6) / NPS (1-6) (Human)
758 Neuropeptide S (1-7) / NPS (1-7) (Human)
759 Neuropeptide S (1-8) / NPS (1-8) (Human)
760 Prepro-Neuropeptide S (24-48) (Rat)
761 Prepro-Neuropeptide S (23-67) (Human)
762 Prepro-Neuropeptide S (23-67) / Prepro-NPS (23-67) (Mouse)
763 Neuroendorcine Regulatory Peptide-1 (NERP-1) (Human)
764 Neuroendorcine Regulatory Peptide-2 (NERP-2) (Human)
765 Prepro -Neurotensin (24-55)
766 Prepro-Neurotensin (115-140)
767 Prepro-Neuropeptide B (56-81) (Human)
768 Copeptin (Human)
769 Prepro-Vasopressin Neurophysin (153-166) (Mouse)
770 Prepro-Vasopressin Neurophysin (130-143) (Mouse)
771 Prepro-Vasopressin Neurophysin (158-168) Mouce)
772 FLP-18 (DFDGAMPGVLRF-NH2)
773 FLP-18 (EMPGVLRF-NH2)
774 Contulakin-G Precursor (51-76) / CGX-1160 (51-76) (*Conus geographus*)
775 Conantokin-G Precursor (81-97) Amide
776 FM059a (GPR-54 agonist)
777 Peptide-34 (GPR-54 agonist)
778 Nesfatin-1 (1-82) (Human)
779 Nesfatin-1 (Rat)
780 Pro-Angiotensin-12 (Rat)
781 Angiotensin A
782 Spexin (Human)
783 Prepro-Spexin (Human)
784 Alarin (Human)
785 Alarin (Mouse)
786 Alarin (Rat)
787 Salusin (40-mer) / Prepro-Salusin (30-69) (Human)
788 Salusin (50-mer) / Salusin-β-RR-Salusin-α (Human)
789 TLQP-21 (Human)
790 TLQP-21 (Rat, Mouse)
791 Neuronostatin-13 Amide (Human, Porcine)
792 Neuronostatin-13 free acid (Human, Porcine)
793 Neuronostatin-13 Amide (Rat, Mouse)
794 Neuronostatin-19 Amide (Human, Porcine)
795 Neuronostatin-19 Amide (Rat, Mouse)
796 Neuronostatin-11 Amide (Human, Porcine)
797 Copeptin (Rat)
798 Hemopressin (Rat, Mouse, Porcine)
799 hemopressin (Human)
800 Oxyntomodulin (Human,Rat, Mouse)
801 Oxyntomodulin (30-37) (Porcine)
802 Oxyntomodulin (32-37) (Human, Rat, Mouse)
803 Oxyntomodulin (19-37) (Human, Rat, Mouse)
804 Oxyntomodulin (30-37) (Human, Rat, Mouse)
805 Glucagon-like peptide -1 (9-36) Amide (Human)
806 Glicentin-related Polypeptide (21-50) (Human)
807 Exendin-4 (1-9)

808 GLP-1 (7-36) Amide, pGlu1 (Human)
809 Ac-GLP-1 (7-36) Amide (Human)
810 Hypocholesterolemic peptide
811 Musclin/OSTN (83-112) (Human)
812 Musclin/OSTN (80-102) (Human)
813 Musclin / OSTN (116-133) (Human)
814 Musclin/OSTN (108-133) (Human)
815 Musclin (113-130) (Mouse) / Musclin (115-132) (Rat)
816 Musclin/OSTN (83-132) (Human)
817 Musclin (105-130) (Mouse)
818 Musclin/OSTN (83-133) (Human)
819 Musclin (80-130) (Mouse)
820 Musclin/OSTN (31-78) (Human)
821 Musclin (39-75) (Mouse)
822 Musclin (31-77) (Rat)
823 CU-NP (T-32-Y), lot number 426283
824 Adrenomedullin -Gly (13-53) (Human)
825 Adrenomedullin-5 (Porcine)
826 Catestatin (Human)
827 Catestatin (Mouse)
828 Catestatin (Rat)
829 Vasostatin I / Prepro-Chromogranin A (19-94) (Human)
830 Vasostatin I (17-43) (Human)
831 Vasostatin I (17-76) (Human)
832 Adiponectin, globular form, (Human)
833 Adiponectin, globular form, (Rat)
834 Anginex
835 FGF-21 (Mouse), recombinant
836 FGF-21 (Human), recombinant
837 FGF-21 (26-47) (Human)
838 FGF-21 (98-124) (Human)
839 FGF-21 (184-209) , Cys0, (Human)
840 Angiotensin II Receptor, Type I (181-187) (Human)
841 Enterostatin (Human, Rat, Mouse)
842 Bombinin-1 like peptide 1
843 Octapeptide 1
844 Octapeptide 2
845 Bombinakinin-GAP
846 Bombinakinin-M
847 Maximakinin
848 KassinaKinin S
849 Prepro-Augurin (42-65) (Human)
850 Prepro-Augurin (71-107) (Human)
851 Prepro-Augurin (108-132) (Human)
852 Prepro Augurin (133-148) (Human)
853 Neuropeptide S, Tyr15, (Mouse)
854 AF-16 Peptide (Human)
855 Sleepless (SSS) (131-158)
856 Sleepless (SSS) (42-63)
857 PYY (2-36)-Gly (Human)
858 PYY (4-36)-Gly (Human)
859 PYY (4-36)-Gly (Rat, Canine)
860 GIP, Pro3, (Rat)
861 GIP (3-42) (Human)

| | |
|---|---|
| 862 | GIP (Mouse) |
| 863 | GIP, (D-Ala2) (Human) |
| 864 | Gastrin I (Porcine) |
| 865 | Xendorphin B 1 |
| 866 | Cortagine |
| 867 | QV-A1 |
| 868 | QV-A2 |
| 869 | QV-A3 |
| 870 | QV-A4 |
| 871 | QV-A5 |
| 872 | QV-A6 |
| 873 | QV-B1 |
| 874 | QV-B2 |
| 875 | QV-B3 |
| 876 | QV-B4 |
| 877 | QV-B5 |
| 878 | QV-B6 |
| 879 | QV-C1 |
| 880 | QV-C2 |
| 881 | QV-C3 |
| 882 | QV-C4 |
| 883 | QV-C5 |
| 884 | QV-C6 |
| 885 | QV-D1 |
| 886 | QV-D2 |
| 887 | QV-D3 |
| 888 | QV-D4 |
| 889 | QV-D5 |
| 890 | QV-D6 |
| 891 | QV-E1 |
| 892 | QV-E2 |
| 893 | QV-E3 |
| 894 | QV-E4 |
| 895 | QV-E5 |
| 896 | QV-E6 |
| 897 | QV-F1 |
| 898 | QV-F2 |
| 899 | QV-F3 |
| 900 | QV-F4 |
| 901 | QV-F5 |
| 902 | QV-F6 |
| 903 | QV-G1 |
| 904 | QV-G2 |
| 905 | QV-G3 |
| 906 | QV-G4 |
| 907 | QV-G5 |
| 908 | QV-G6 |
| 909 | QV-H1 |
| 910 | QV-H2 |
| 911 | QV-H3 |
| 912 | QV-H4 |
| 913 | QV-H5 |
| 914 | QV-H6 |
| 915 | QV-A7 |

916 QV-A8
917 QV-A9
918 QV-A10
919 QV-A11
920 QV-A12
921 QV-B7
922 QV-B8
923 QV-B9
924 QV-B10
925 QV-B11
926 QV-B12
927 QV-C7
928 QV-C8
929 QV-C9
930 QV-C10
931 QV-C11
932 QV-C12
933 QV-D7
934 QV-D8
935 QV-D9
936 QV-D10
937 QV-D11
938 QV-D12
939 QV-E7
940 QV-E8
941 QV-E9
942 QV-E10
943 QV-E11
944 QV-E12
945 QV-F7
946 QV-F8
947 QV-F9
948 QV-F10
949 QV-F11
950 QV-F12
951 QV-G7
952 QV-G8
953 QV-G9
954 QV-G10
955 QV-G11
956 QV-G12
957 Chemerin9 peptides&elephants, custom synthesis
958 KE108 750nM Bachem # H-6276
959 Neuropeptide S 750nM Bachem # H-6162
960 Apelin12 750nM peptides&elephants, custom synthesis
961 BMAP-27 Phoenix Pharmaceuticals # 012-29
962 Neurotensin peptides&elephants, custom synthesis
963 GLP2-Gly2 peptides&elephants, custom synthesis
964 KE108 Bachem # H-6276
965 Neuropeptide S Bachem # H-6162
966 Apelin12 peptides&elephants, custom synthesis
967 P-58-4-Gly Amide Phoenix Pharmaceuticals # 036-30
968 KE108 Bachem # H-6276
969 Lanreotide 750nM Bachem # H-9055

970 Gonadoliberin 750nM Bachem # H-4005
971 Urotensin II 750nM peptides&elephants, custom synthesis
972 VIP peptides&elephants, custom synthesis
973 Substance P Bachem # H-1890
974 Bradykinin peptides&elephants, custom synthesis
975 Lanreotide Bachem # H-9055
976 Gonadoliberin Bachem # H-4005
977 Urotensin II ppeptides&elephants, custom synthesis
978 Xenopsin Bachem # H-4885
979 Apelin12 peptides&elephants, custom synthesis
980 ATP Sigma # A7699
981 Ghrelin 750nM peptides&elephants, custom synthesis
982 aEndorphin 750nM Bachem # H-2695
983 Angiotensin Converting Enzyme Inhibitor / SQ20991 Phoenix Pharmaceuticals #002-38
984 aEndorphin Bachem # H-2695
985 Carbachol Sigma # C4382
986 Somatostatin14 peptides&elephants, custom synthesis
987 Ghrelin peptides&elephants, custom synthesis
988 aEndorphin Bachem # H-2695
989 ßEndorphin Bachem # H-2700
990 Octreotide peptides&elephants, custom synthesis
991 Apamin 750nM Sigma # A9459
992 GIP-750nM peptides&elephants, custom synthesis
993 GLP1 750nM Bachem # H-5102
994 Chemerin9 750nM peptides&elephants, custom synthesis
995 Motilin 750nM Bachem # H-4385
996 Neurotensin 750nM peptides&elephants, custom synthesis
997 GIP1-30 peptides&elephants, custom synthesis
998 GLP1 Bachem # H-5102

**Supplementary table 2: Confirmed hits from screening**

| # | ligand | DMR assay | calcium assay |
|---|---|---|---|
| 1 | Adrenomedullin (ADM), (16-52) (Human) | + | - |
| 2 | Alpha Endorphin (Beta-Lipotropin (61-76)) | - | + |
| 3 | Alpha-Melanocyte Stimulating Hormone | - | + |
| 4 | Angiotensin A | + | + |
| 5 | Angiotensin I (1-9) (Human) | + | - |
| 6 | Angiotensin I (Human) | + | + |
| 7 | Angiotensin I, [Des-Asp1-] (Human) | + | + |
| 8 | Angiotensin II (Human) | + | + |
| 9 | Angiotensin II [Sar1] | + | + |
| 10 | Angiotensin II, [Des-Asp1-] (Angiotensin III) (Human) | + | + |
| 11 | Angiotensin IV [AT-II (3-8)] | - | + |
| 12 | Atriopeptin II (Rat Rabbit Mouse) | - | + |
| 13 | Lanreotide (peptides&elephants) 750 nM | + | - |
| 14 | Lanreotide (peptides&elephants) 75 nM | + | - |
| 15 | Bombesin | - | + |
| 16 | Bombesin (6-14), [D-Phe6, Beta-Ala11, Phe13, Nle14] | - | + |
| 17 | Bradykinin | + | + |
| 18 | Bradykinin [Ile-Ser-] (T-Kinin) | + | + |
| 19 | Bradykinin (BACHEM) | + | + |
| 20 | Calcitonin (Eel) | + | - |
| 21 | Calcitonin (Human) | + | - |
| 22 | Calcitonin (Rat) | + | - |
| 23 | Calcitonin (salmon) | + | - |
| 24 | Calcitonin Gene Related Peptide (CGRP) (Human) | + | - |
| 25 | Calcitonin Gene Related Peptide (CGRP) (8-37) (Rat) | + | - |
| 26 | Calcitonin Gene Related Peptide (CGRP) (Rat) | + | - |
| 27 | Calcitonin Gene Related Peptide II (CGRP II) (Human) | + | - |
| 28 | Calcitonin II (Salmon) | + | - |
| 29 | Calcitonin III (salmon) | + | - |
| 30 | Calcitonin IV (salmon) | + | - |
| 31 | Calcitonin V (salmon) | + | - |
| 32 | Endorphin, Beta-Neo | + | + |
| 33 | Gastrin Releasing Peptide (GRP) (14-27) (Porcine, Human) | - | + |
| 34 | Neuromedin C (Porcine) | - | + |
| 35 | Pro-Angiotensin-12 (Rat) | + | + |
| 36 | RFRP-1 (Human) | + | - |
| 37 | Somatostatin-25 | + | - |
| 38 | Somatostatin-28 | + | - |

**Supplementary Table S3 | Clinical characteristics of the tissues analyzed by receptor autoradiography.**

| # | Sex | Age at surgery | Tissue type | Primary | Origin of tissue | Ki67 | Grade |
|---|---|---|---|---|---|---|---|
| **1** | m | 75 | primary | pancreas | pancreas | 3 % | G2 |
| **2** | m | 67 | metastasis | pancreas | liver | 2 % | G1 |
| **3** | m | 28 | primary | pancreas | pancreas | 6 % | G2 |
| **4** | f | 43 | metastasis | pancreas | liver | 5 % | G2 |
| **5** | f | 44 | metastasis | pancreas | liver | 25 % | G3 |
| **6** | f | 51 | primary | pancreas | pancreas | 1 % | G1 |
| **7** | f | 38 | metastasis | pancreas | liver | 80 % | G3 |
| **8** | f | 65 | primary | pancreas | pancreas | 15 % | G2 |
| **9** | m | 34 | primary | ileum | ileum | 10 % | G2 |
| **10** | m | 74 | metastasis | ileum | liver | 3 % | G2 |
| **11** | f | 54 | metastasis | ileum | liver | 5 % | G2 |
| **12** | m | 76 | metastasis | ileum | liver | 2 % | G1 |
| **13** | f | 54 | primary | ileum | ileum | 5 % | G2 |
| **14** | m | 49 | primary | ileum | ileum | 4 % | G2 |
| **15** | f | 57 | metastasis | ileum | liver | 5 % | G2 |
| **16** | f | 62 | primary | ileum | ileum | 1-2 % | G1 |
| **17** | f | 53 | primary | ileum | ileum | 1 % | G1 |
| **18** | m | 58 | normal | | pancreas | | |
| **19** | m | 67 | normal | | pancreas | | |
| **20** | m | 50 | normal | | pancreas | | |
| **21** | f | 65 | normal | | pancreas | | |
| **22** | m | 65 | normal | | pancreas | | |
| **23** | m | 34 | normal | | ileum | | |
| **24** | f | 65 | normal | | duodenum | | |
| **25** | m | 72 | normal | | lymph node ileum | | |
| **26** | f | 69 | normal | | ileum | | |
| **27** | m | 65 | normal | | duodenum | | |
| **28** | m | 50 | normal | | liver | | |

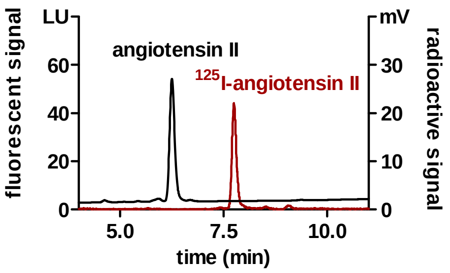


**Supplementary Figure S1: HPLC purification of radiolabeled angiotensin II.** For radioligand binding experiments angiotensin II was labeled with iodine-125 and purified from unlabeled peptide by HPLC. Chromatogram showing peptide peaks as recorded by fluorescence detector (black line, ex=280 nM, em=340 nm) and radioactive detector (red line) with a clear difference of retention times.

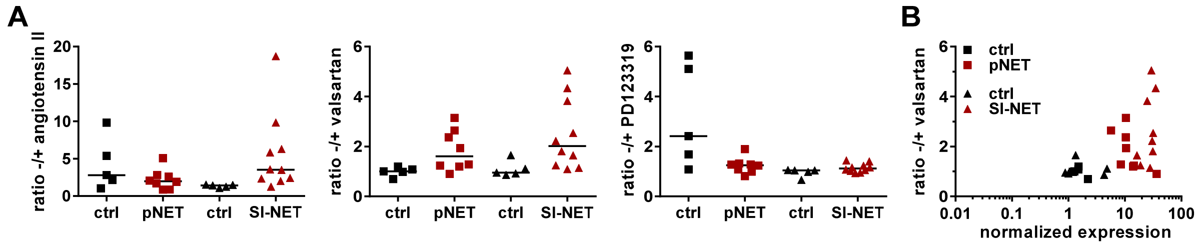


**Supplementary Figure S2: Quantification of autoradiography by an alternative method.** In addition to ImageJ analysis, sections were wiped off after film development and the counts per minute (cpm) measured with a gamma counter. Ratios of total to non-specific binding are shown for angiotensin II, valsartan and PD123319 **(A)**. **(B)** Scatter plot showing the correlation of AGTR1 gene expression (x-axis, RT-qPCR) with receptor protein expression (y-axis, autoradiography). ctrl, control; pNET, pancreatic NET; SI-NET, small-intestinal NET.

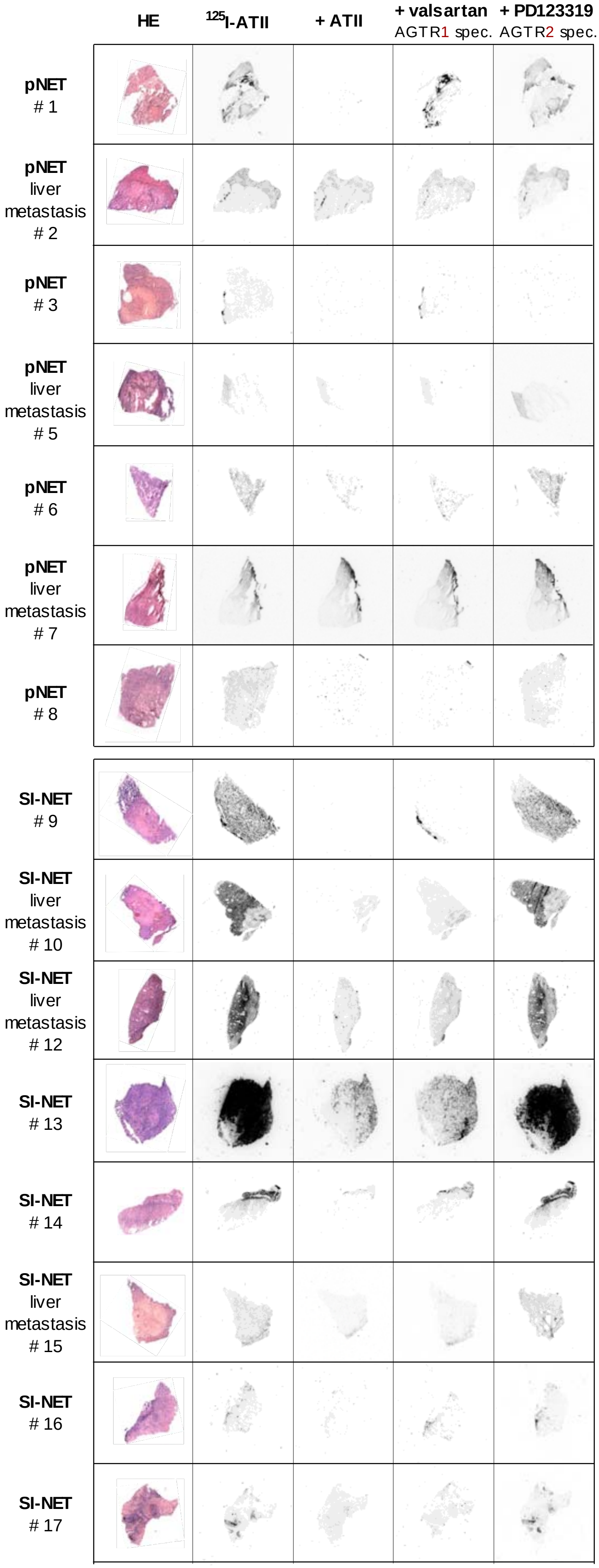


**Supplementary Figure S3: In vitro receptor autoradiography of pancreatic and small-intestinal NET tissues.** For each pancreatic (pNET) and small-intestinal (SI-NET) tumor tissue, HE staining and autoradiograms of adjacent cryosections are displayed. Cryosections were either incubated with $^{125}$I-angiotensin II alone ($^{125}$I-ATII, total binding) or in the presence of additional 1 µM unlabeled angiotensin II (ATII), AGTR1 antagonist valsartan or AGTR2 antagonist PD123319 (non-specific binding).

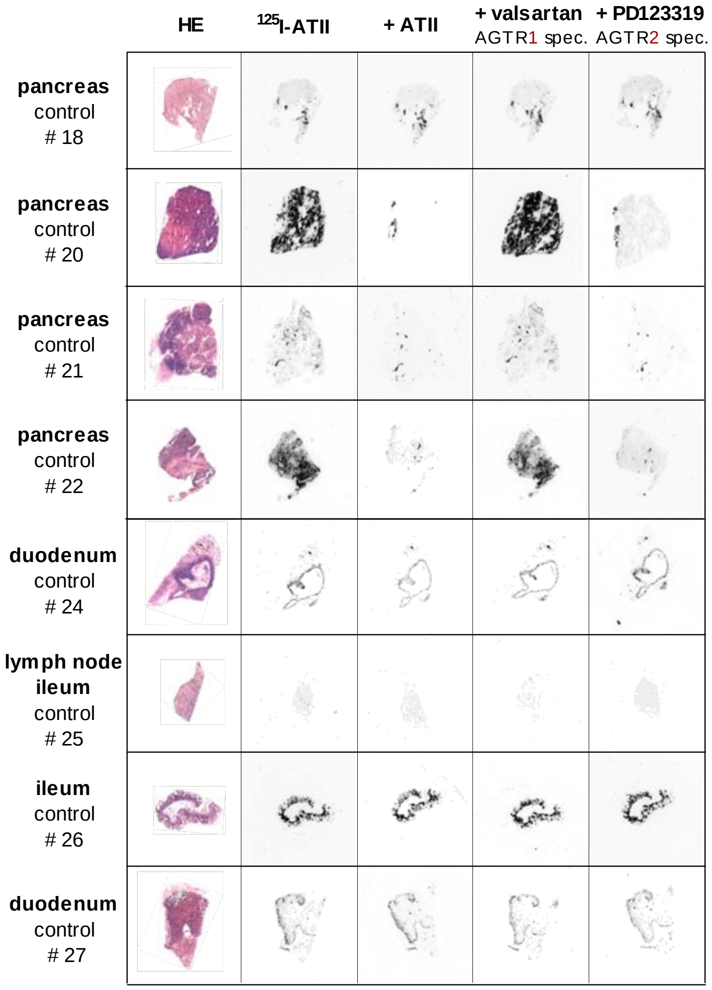


**Supplementary Figure S4: In vitro receptor autoradiography of pancreatic and small-intestinal control tissues.** For each tissue, HE staining and autoradiograms of adjacent cryosections are displayed. Cryosections were either incubated with $^{125}$I-angiotensin II alone ($^{125}$I-ATII, total binding) or in the presence of additional 1 µM unlabeled angiotensin II (ATII), AGTR1 antagonist valsartan or AGTR2 antagonist PD123319 (non-specific binding).

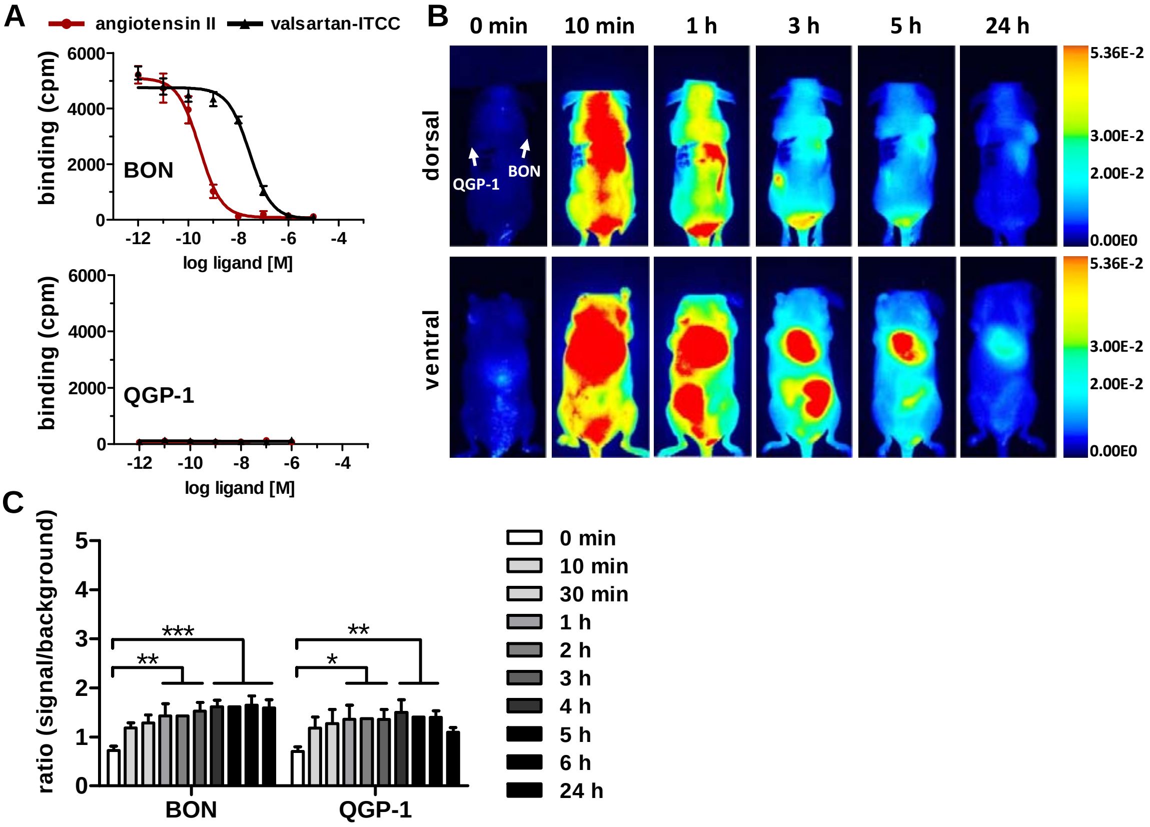


**Supplemetary Figure S5: In vivo near-infrared fluorescent (NIRF) imaging with AGTR1-targeting valsartan-ITCC. (A)** AGTR1-positive BON and AGTR1-negative QGP-1 cells were incubated with $^{125}$I-angiotensin II and increasing concentrations of unlabeled angiotensin II or valsartan-ITCC. Data show mean ± S.E.M. for BON (n=3-4) or mean ± S.D. for QGP-1 (n=1). **(B)** Biodistribution of 1 nmol i.v. valsartan-ITCC in a mouse model subcutaneously injected with BON (right shoulder) and QGP-1 (left shoulder). Images were acquired at the indicated time points before or after injection and are displayed with an equally adjusted gain for either dorsal or ventral signals. **(C)** Signals from in vivo NIRF imaging were quantified by calculating the ratio of signal (tumor) to background (neck) for valsartan-ITCC. Bars represent mean ± S.E.M. of three different animals. Evaluated with matched two-way ANOVA and Bonferroni post-hoc test, * P<0.05, ** P<0.01, *** P<0.001.

### Supplementary Methods

#### Synthesis of valsartan-ITCC

Valsartan-ITCC (the dye ITCC linked via 1,3-diamino propane) was synthesized as follows.

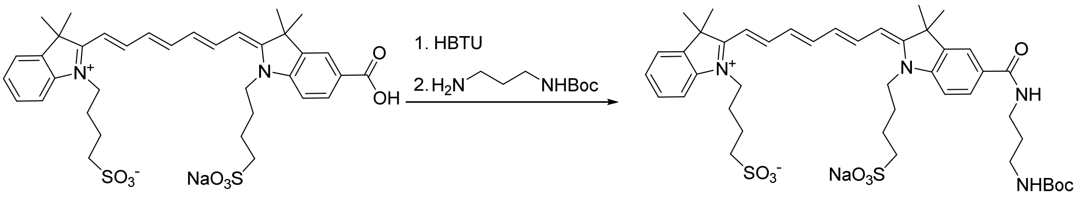


The ITCC dye (210 mg, 278 µmol, 1.0 eq.) was dissolved with HBTU (179 mg, 473 µmol, 1.7 eq.) in DMF (5.7 mL). DIPEA (174 µL, 1.02 mmol, 3.7 eq.) was added and the reaction mixture stirred for 1.5 h. *N*-Boc-1,3-diamino propane (72.7 mg, 417 µmol, 1.5 eq.) and DIPEA (87 µL, 510 µmol, 1.8 eq.) were dissolved in DMF (1.9 mL) and added to the reaction mixture which was stirred for additional 15 h. The mixture was precipitated from $Et_2O$ and the precipitate purified via flash chromatography (DCM/MeOH) and reverse phase column chromatography ($H_2O$/MeOH) to obtain the product as a green solid (69 mg). MS (ESI): $C_{44}H_{59}N_4NaO_9S_2$ $[M+Na]^+$ calculated: 897.3513; found: 897.3469.

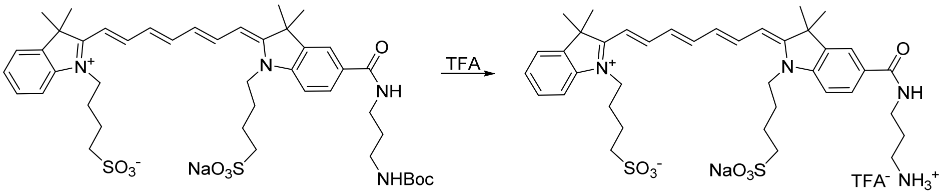


The Boc-protected ITCC dye (69 mg) was dissolved in DCM (3.0 mL) and subjected to deprotection by addition of TFA (300 µL). The reaction mixture was stirred for 2 h, the solvent removed under reduced pressure and the mixture codistilled with DCM several times to remove TFA. After filtration with methanol, a green solid was obtained (41 mg). MS (ESI): $C_{39}H_{51}N_4NaO_7S_2$ $[M+Na]^+$ calculated: 797.2989; found: 797.2967.

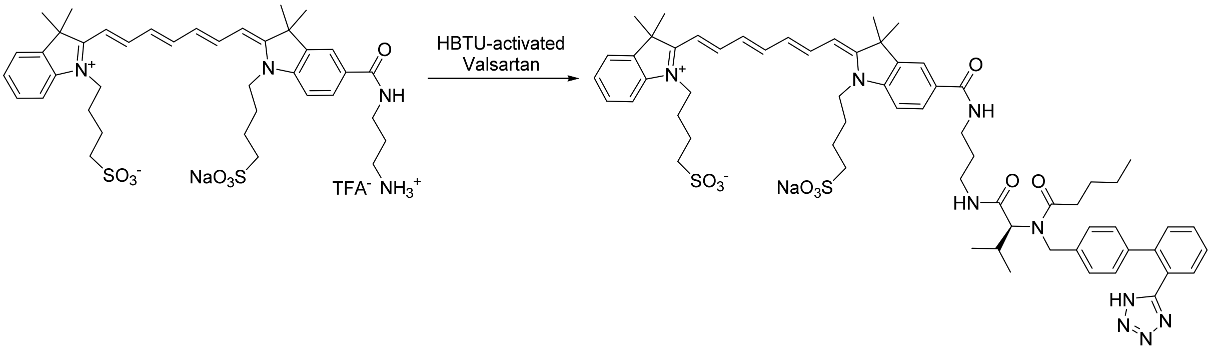


Valsartan (5.0 mg, 12 µmol, 1.3 eq.) was dissolved in DMF (125 µL) with HBTU (7.4 mg, 20 µmol, 2.2 eq.) and DIPEA (7.2 µL, 42 µmol, 4.7 eq.) and the reaction mixture stirred for 1 h. The deprotected ITCC dye (8.0 mg, 9.0 µmol, 1.0 eq.) was dissolved in DMF (125 µL) with DIPEA (3.6 µL, 21 µmol, 2.3 eq.) and added to the reaction mixture which was stirred for additional 17 h. The mixture was precipitated from $Et_2O$ and the precipitate purified via flash chromatography (DCM/MeOH) to afford the product as a green solid (3.9 mg).MS (ESI): $C_{63}H_{78}N_9NaO_9S_2$ $[M]^{\cdot+}$ calculated: 1191.5262; found: 1191.5257.
